# Differential usage of DNA modifications in neurons, astrocytes, and microglia

**DOI:** 10.1101/2023.06.05.543497

**Authors:** Kyla B. Tooley, Ana J. Chucair-Elliott, Sarah R. Ocañas, Adeline H. Machalinski, Kevin D. Pham, David R. Stanford, Willard M. Freeman

**Affiliations:** Department of Physiology, University of Oklahoma Health Sciences Center, Oklahoma City, OK USA; Genes & Human Disease Program, Oklahoma Medical Research Foundation, Oklahoma City, OK USA; Center for Biomedical Data Sciences, Oklahoma Medical Research Foundation, Oklahoma City, OK USA; Department of Biochemistry, University of Oklahoma Health Sciences Center, Oklahoma City, OK USA; Oklahoma City Veterans Affairs Medical Center, Oklahoma City, OK USA

**Keywords:** Brain, epigenomics, methylation, hydroxymethylation, regulatory elements, genome regulation

## Abstract

**Background:** Cellular identity is determined partly by cell type-specific epigenomic profiles that regulate gene expression. In neuroscience, there is a pressing need to isolate and characterize the epigenomes of specific CNS cell types in health and disease. This is especially true as for DNA modifications where most data are derived from bisulfite sequencing that cannot differentiate between DNA methylation and hydroxymethylation. In this study, we developed an *in vivo* tagging mouse model (Camk2a-NuTRAP) for paired isolation of neuronal DNA and RNA without cell sorting and then used this model to assess epigenomic regulation of gene expression between neurons and glia.

**Results:** After validating the cell-specificity of the Camk2a-NuTRAP model, we performed TRAP-RNA-Seq and INTACT whole genome oxidative bisulfite sequencing to assess the neuronal translatome and epigenome in the hippocampus of young mice (3 months old). These data were then compared to microglial and astrocytic data from NuTRAP models. When comparing the different cell types, microglia had the highest global mCG levels followed by astrocytes and then neurons, with the opposite pattern observed for hmCG and mCH. Differentially modified regions between cell types were predominantly found within gene bodies and distal intergenic regions, with limited differences occurring within proximal promoters. Across cell types there was a negative correlation between DNA modifications (mCG, mCH, hmCG) and gene expression at proximal promoters. In contrast, a negative correlation of mCG with gene expression within the gene body while a positive relationship between distal promoter and gene body hmCG and gene expression was observed. Furthermore, we identified a neuron-specific inverse relationship between mCH and gene expression across promoter and gene body regions.

**Conclusions:** In this study, we identified differential usage of DNA modifications across CNS cell types, and assessed the relationship between DNA modifications and gene expression in neurons and glia. Despite having different global levels, the general modification-gene expression relationship was conserved across cell types. The enrichment of differential modifications in gene bodies and distal regulatory elements, but not proximal promoters, across cell types highlights epigenomic patterning in these regions as potentially greater determinants of cell identity.

## Background

DNA methylation (mC) and hydroxymethylation (hmC) are stable modifications added to the 5’ position of the cytosine ring in the CpG (CG) and non-CpG (CH) contexts, each (mCG, hmCG, and mCH) having distinct roles in genome regulation and gene expression in the central nervous system (CNS) [1]. The presence of hmCH is debated, and it’s potential role in genome regulation has yet to be elucidated [2-4]. DNA modification patterns modulate CNS cell differentiation and specialization [5-8], with deposition and removal occurring at different points of neurodevelopment [2]. While mCG has been extensively studied, there is increasing interest in investigating hmCG and mCH in neuroscience research due to their higher abundance in the brain compared to other tissues [9-11] and their potential involvement in neurological disease [12-16]. Notably, the deposition of mCH coincides with increased synaptic density and a positive association between gene body hmCG and gene expression suggests potential functional roles of DNA modifications in both neurodevelopment and the adult brain [2, 17].

The complete role of DNA modifications in regulating gene expression is still being determined, but recent advances have revealed that different modifications have distinct relationships to gene expression that can vary by genomic context. mCG exerts a well-established repressive function on gene expression when deposited in the proximal promoter region [18]. Methyl-binding proteins, such as MeCP2, recognize and bind to methylated DNA, further impeding transcription and reinforcing the repressive effect of mCG [19]. This repressive function can have long-lasting effects, as mCG plays a crucial role in the long-term repression of repetitive elements and X-chromosome inactivation within the CNS [10, 11]. Within gene bodies, mCG has been described exhibiting both a negative [3, 20] and positive [21] relation to gene expression, leaving the functional relationship of this modification to gene expression somewhat ambiguous. In contrast, hmCG is positively correlated with gene expression and is enriched at tissue-specific genes and transcription factor binding sites [22, 23]. In postmitotic neurons, hmCG is primarily located in the gene body of expressed genes, and is attributed to “functional demethylation” of these regions, serving to decrease binding affinity of MeCP2 and promote gene expression [24, 25]. Furthermore, it is hypothesized that hmCG may be required for development of the complex morphology and synaptic connections of long-range postmitotic neurons [26, 27].

Methylation in CH contexts (C followed by C, A, or T) was not previously considered a prominent site of cytosine methylation. However, neurons have the highest proportion of mCH in the brain [2]. mCH is depleted within highly expressed genes and their regulatory elements, instead serving to fine tune the cell type-specific expression of lowly expressed neuronal genes [6, 28]. Additionally, mCH accumulation during development parallels synaptogenesis, indicating that mCH is likely important in regulating the formation and maintenance of synaptic connections [29].

Although the neuro-epigenomics research has advanced considerably, the field lacks comprehensive maps of the relationships between DNA modifications (mCG, hmCG, mCH) and gene expression within neurons and glia. To address the technical and knowledge gaps in the field we combine the <u>Nu</u>clear Tagging and <u>T</u>ranslating <u>R</u>ibosome <u>A</u>ffinity <u>P</u>urification (NuTRAP) mouse line [30-32] with a well-established neuronal-specific inducible cre-recombinase system (Camk2a-cre/ERT2 [33-36]) to perform a paired translatomic and epigenomic analysis of excitatory glutamatergic pyramidal neurons in the hippocampus [37-40]. To gain a broader perspective, we compare our neuronal findings to astrocytic and microglial data [41]. This comparative analysis reveals cell-type specific usage of DNA modifications and their associations with mRNA expression across three CNS cell types. Studies described here: 1) validate the Camk2a-NuTRAP model, 2) compare DNA modification usage across three CNS cell types, and 3) assess the relationship between DNA modifications and mRNA levels in the three CNS cell types, providing insight into the regulatory mechanisms governing gene expression. By undertaking these investigations, we hope to advance the understanding of the role of DNA modifications in gene regulation across different CNS cell types, paving the way for future discoveries in neuro-epigenomics.

## Results

### Immunohistochemical validation of the Camk2a-NuTRAP mouse brain

To ensure minimal interference with neurodevelopmental processes, we performed tamoxifen (Tam) induction of Camk2a-NuTRAP in mature adult mice at 3 months of age (3mo). This timing was chosen to avoid deficits in spatial learning, contextual fear memory, and presynaptic structure, that can arise after perturbing Camk2a expression during early neurodevelopment [42, 43]. Brains were collected one month following Tam induction and sectioned sagittally for immunohistochemical analysis.

Immunostaining of Camk2a-NuTRAP (Camk2a-cre/ERT2^+^; NuTRAP^+^) brains showed EGFP and mCherry colocalization in cells expressing the pan-neuronal marker NeuN. No EGFP or mCherry expression was seen in Camk2a-cre negative counterparts (**Figure 1A**) and minimal expression was observed in a portion of NeuN^+^ cells of Camk2a-NuTRAP (-Tam) brains, which is consistent with previous reports (**Supplemental Figure 1B**) [36, 44]. Camk2a-NuTRAP (+Tam) brains show no expression of EGFP in microglial (CD11b^+^), endothelial (CD31^+^), or astrocytic (GFAP^+^) cells (**Supplemental Figure 1C-E**). Collectively, these findings indicate a robust neuronal-specific and tamoxifen-dependent induction of the NuTRAP allele. This validation ensures that the experimental manipulations specifically target neuronal cells while minimizing any confounding effects on other cell types in the brain.

**Figure 1:**
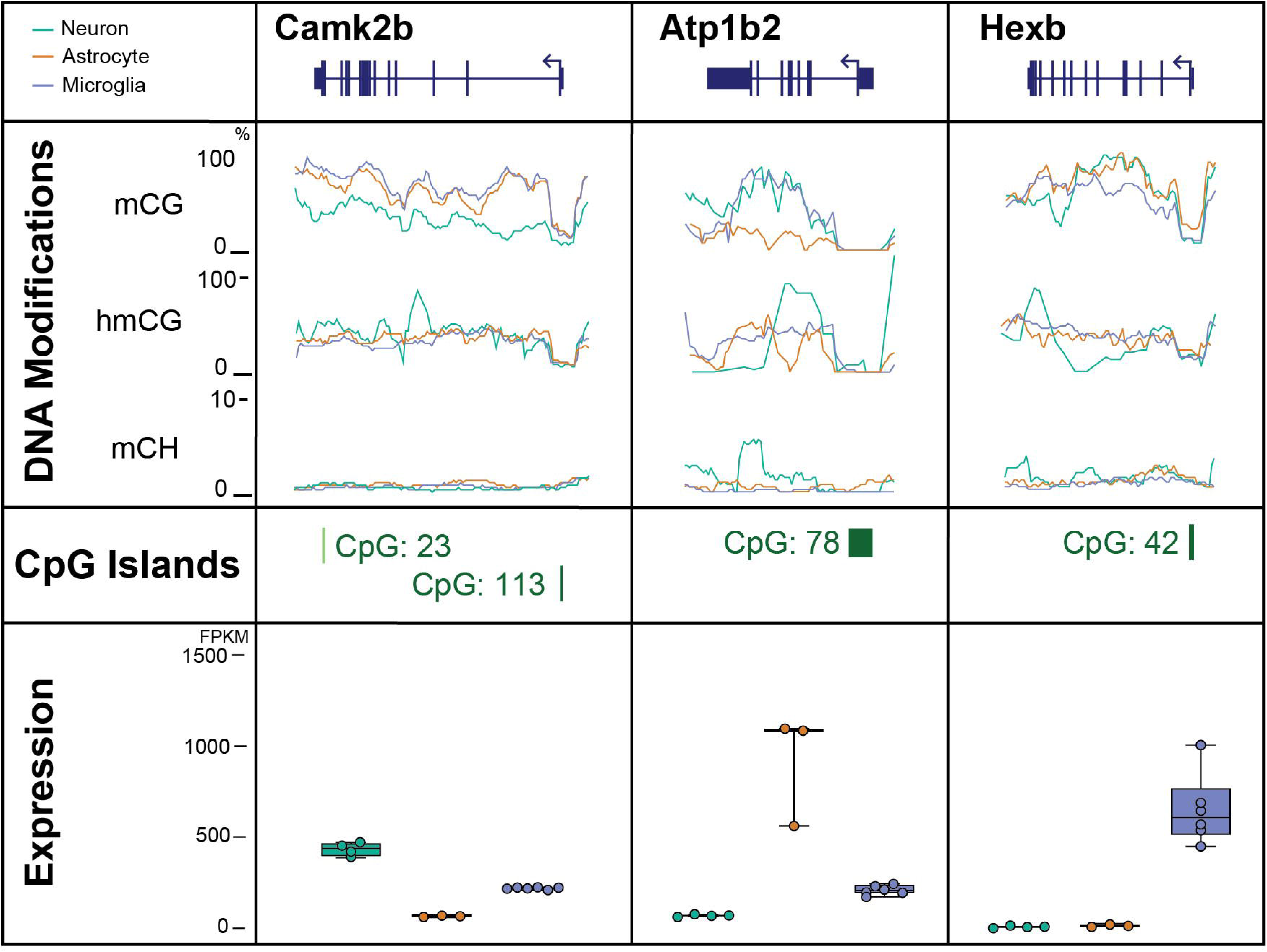
Validation of neuronal translatome enrichment in TRAP-RNA from Camk2a-NuTRAP mouse hippocampus. **A)** Imaging of the hippocampal dentate gyrus demonstrated EGFP and mCherry co-expression with NeuN+ cells in Camk2a-cre^+^; NuTRAP^+^ brains, but not Camk2a-cre^-^; NuTRAP^+^ brains. **B)** TRAP-isolated hippocampal RNA from input, negative, and positive fractions were assessed by qPCR for enrichment and depletion of canonical marker genes for microglia, astrocytes, oligodendrocytes, and neurons. Mean relative gene expression ± SEM scaled to input for each gene. *p < 0.05, **p < 0.01, ***p < 0.001, ****p < 0.0001 by RM one-way ANOVA with Tukey’s multiple comparison test across fractions (n=4/group). **C)** RNA-seq was performed for all fractions (n=4/group). Principal component analysis shows separation of the positive from input and negative fraction samples in the first component. **D)** Cell-type marker gene lists were examined for fold change (Positive/Input) enrichment or depletion shows enrichment of neuronal markers and depletion of other cell-type markers in the positive fraction. **E)** CIBERSORTx calculation of cell type composition of each fraction. The positive fraction is estimated to contain 100% neurons. **F)** Genes with significant enrichment (2111) or depletion (2897) in the positive fraction compared to input were identified (FC > |1.25|, p < 0.05, Benjamini Hochberg multiple testing corrections). **G-J)** Gene Ontology enrichment analysis and Ingenuity Pathway Analysis performed on significantly upregulated or downregulated genes (Positive fraction/Input) identified in E.

### Validation of neuronal translatome enrichment from TRAP-isolated RNA

Translating RNA was isolated from the hippocampus of Camk2a-NuTRAP mice via the TRAP method (<u>T</u>ranslating <u>R</u>ibosome <u>A</u>ffinity <u>P</u>urification) [41]. Subsequent RT-qPCR of RNA from the input, negative, and positive TRAP fractions showed a significant enrichment of neuronal marker genes (*Camk2a*, *Hpca*, *Caln1*, *Kcnip4*, *Stmn2*, and *Snap25*) in the positive fraction compared to input and negative fraction. Conversely, there was a depletion of microglial (*Cx3cr1* and *Itgam*), astrocytic (*Aldhl1l* and *Gfap*), and oligodendrocytic (*Mog*) marker genes in the positive fraction as compared to the input and negative fraction (**Figure 1B; Additional file 1B**).

To further characterize the neuronal translatomic profile, TRAP-isolated RNA was subjected to RNA-Seq, and Principal Component Analysis (PCA) was performed. The PCA revealed clear separation of the positive fraction from input and negative fraction in the first component (**Figure 1C**). In order to validate the TRAP-enrichment of neuronal genes and depletion of other cell type-specific genes, we used marker lists generated from previous cell sorting studies [41] (**Additional file 2**). Enrichment of neuronal genes and depletion of astrocytic, microglial, oligodendrocytic, and endothelial genes were evident in the positive fraction as compared to input (**Figure 1D; Additional file 3A**). Notably, there was a high fold depletion of markers for minority cell types like glia and smaller fold-change enrichment for neurons, which make up the majority of the input.

To estimate the cell type composition of the input, negative and positive fractions, we employed CIBERSORTx [45] using established cell type marker lists [41]. This analysis revealed that the input contained the expected variety of cell types at the expected proportions (astrocytes, microglia, neurons, oligodendrocytes, and endothelial cells). The negative fraction demonstrated a depletion of neuronal cells, whereas the positive fraction was estimated to be entirely represented by neurons (∼100%) (**Figure 1E; Additional file 3B**).

Ingenuity Pathway Analysis and Gene Ontology analysis of the significantly enriched and depleted genes in the positive fraction vs input (**Figure 1F; Additional file 3C,D**) revealed enriched genes regulating excitatory neuronal biological processes and functions such as those involved in synaptic structure, maintenance and plasticity (**Figure 1G, H; Additional file 3E,F**). On the other hand, depleted genes were involved in lipid metabolism, immune response, and vascular formation and maintenance, indicating a depletion of genes involved in non-neuronal pathways (**Figure 1I, J; Additional file 3G,H**). Moreover, the positive fraction enrichment of genes involved in spatial learning, a major function of hippocampal neurons, further demonstrated the specificity and relevance of the model in capturing neuronal-specific transcripts [46, 47]. These findings provide valuable insights into the enriched and depleted gene sets within the neuronal translatome, shedding light on the functional processes and pathways associated with neuronal identity and function of the hippocampus.

### Validation of neuronal gDNA isolation from INTACT Whole Genome Bisulfite Sequencing

To ensure the purity of the positive fraction obtained through INTACT isolation (Isolation of <u>N</u>uclei in <u>TA</u>gged in specific <u>C</u>ell <u>T</u>ypes), expression of EGFP within the nucleus [32] was assessed by confocal microscopy. EGFP-positive nuclei surrounded by streptavidin beads were observed in the positive fraction (**Figure 2A**). In contrast, the input showed a mixture of EGFP-positive and EGFP-negative nuclei (**Figure 2B**), while the negative fraction exhibited no EGFP expression (**Figure 2C**).

**Figure 2:**
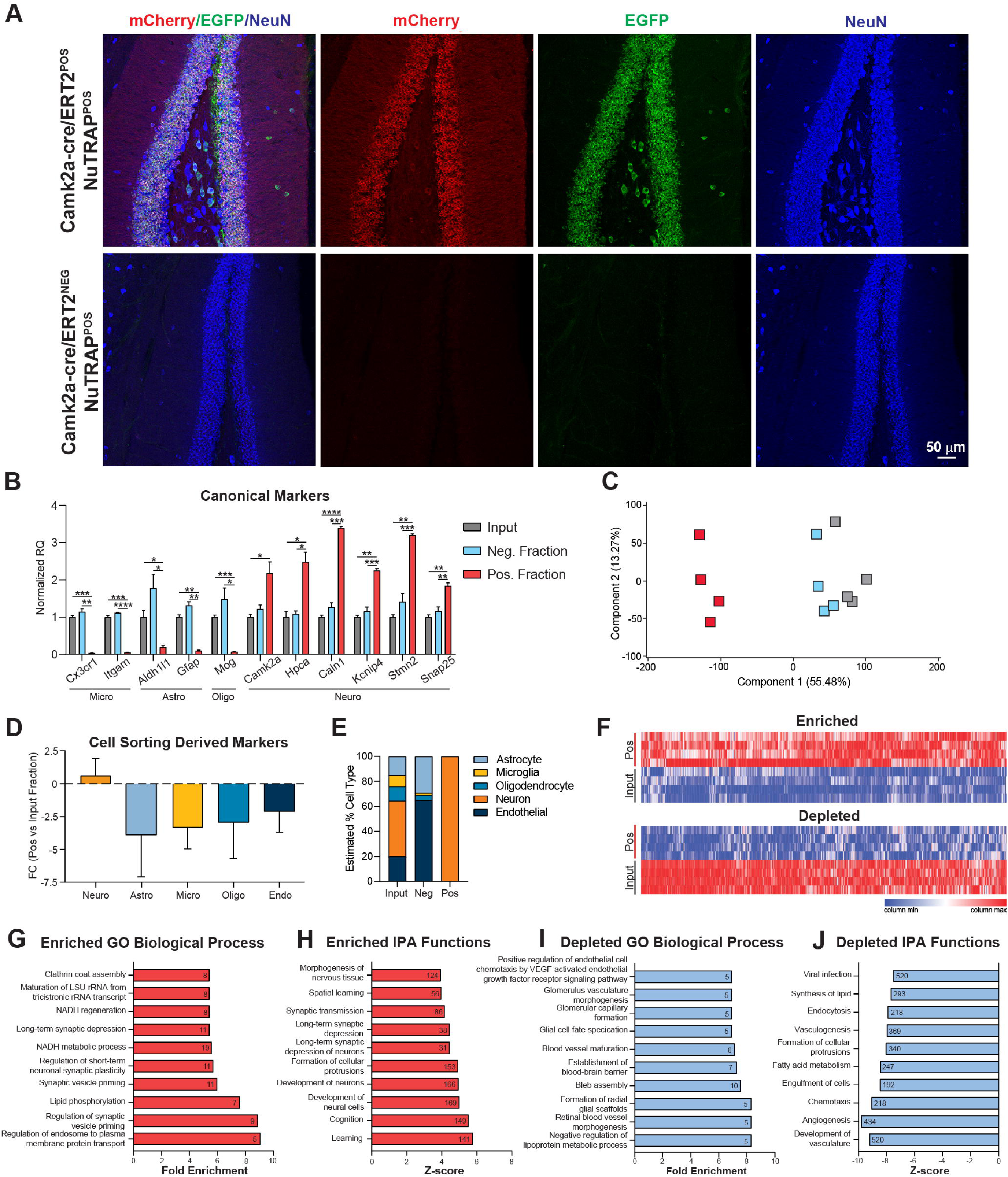
Validation of neuronal genome enrichment in Camk2a-NuTRAP mouse hippocampus by INTACT-BS seq. **A-C)** Confocal images from positive, input, and negative INTACT nuclei isolation fractions demonstrated EGFP+ nuclei in the positive fraction. **D)** INTACT-isolated hippocampal gDNA was bisulfite converted and whole genome levels of CG modifications measured for input, negative, and positive fractions. CG modifications from previously published neuronal methylation studies (hippocampus and cortex) and isolation techniques (Camk2a INTACT, NeuN^+^ sorting, and single cell) were compared to Camk2a-NuTRAP CG modifications. **E)** Whole genome CH modifications were measured for input, negative, and positive fractions. CH modifications from the same neuronal methylation studies from D were compared to Camk2a-NuTRAP CH modifications. **p < 0.01 by one-way ANOVA with Tukey’s multiple testing correction.

To assess DNA modifications in the positive fraction, whole genome oxidative bisulfite sequencing (WGoxBS) was performed on INTACT-isolated gDNA from the input, negative, and positive fractions to measure mC and hmC in the CG and CH contexts. First, the bisulfite-only arm, which detects a combined signal of mC and hmC (total modifications), was compared to previously published neuronal bisulfite sequencing data. Total CG and CH modification levels from the positive fraction were similar to previously published neuronal bisulfite sequencing modification studies [2, 48-50] **(Figure 2D-E)**. These findings provide evidence for the isolation of neuronal-specific genomic DNA, supporting the validity of the INTACT isolation method and the subsequent analysis of DNA modifications in the positive fraction.

### Neuronal epigenome analysis using Whole Genome Oxidative Bisulfite Sequencing

To distinguish between mC and hmC, both whole genome bisulfite sequencing (BS-Seq) and oxidative bisulfite sequencing (oxBS-Seq) were performed on INTACT-isolated DNA from the input, negative and positive fractions (**Supplemental Figure 2A)**. Conversion efficiency, measured by spike-in controls, was close to 100% with no significant variance between samples or groups (**Supplemental Figure 2B-C**). Comparing the different fractions, the positive fraction exhibited significantly lower mCG and higher levels of hmCG and mCH when compared to input and negative fractions (**Figure 3A-C**). Non-CG hydroxymethylation (hmCH) was detected at low levels near background (<1%) and was not significantly different between fractions (**Figure 3D**).

**Figure 3:**
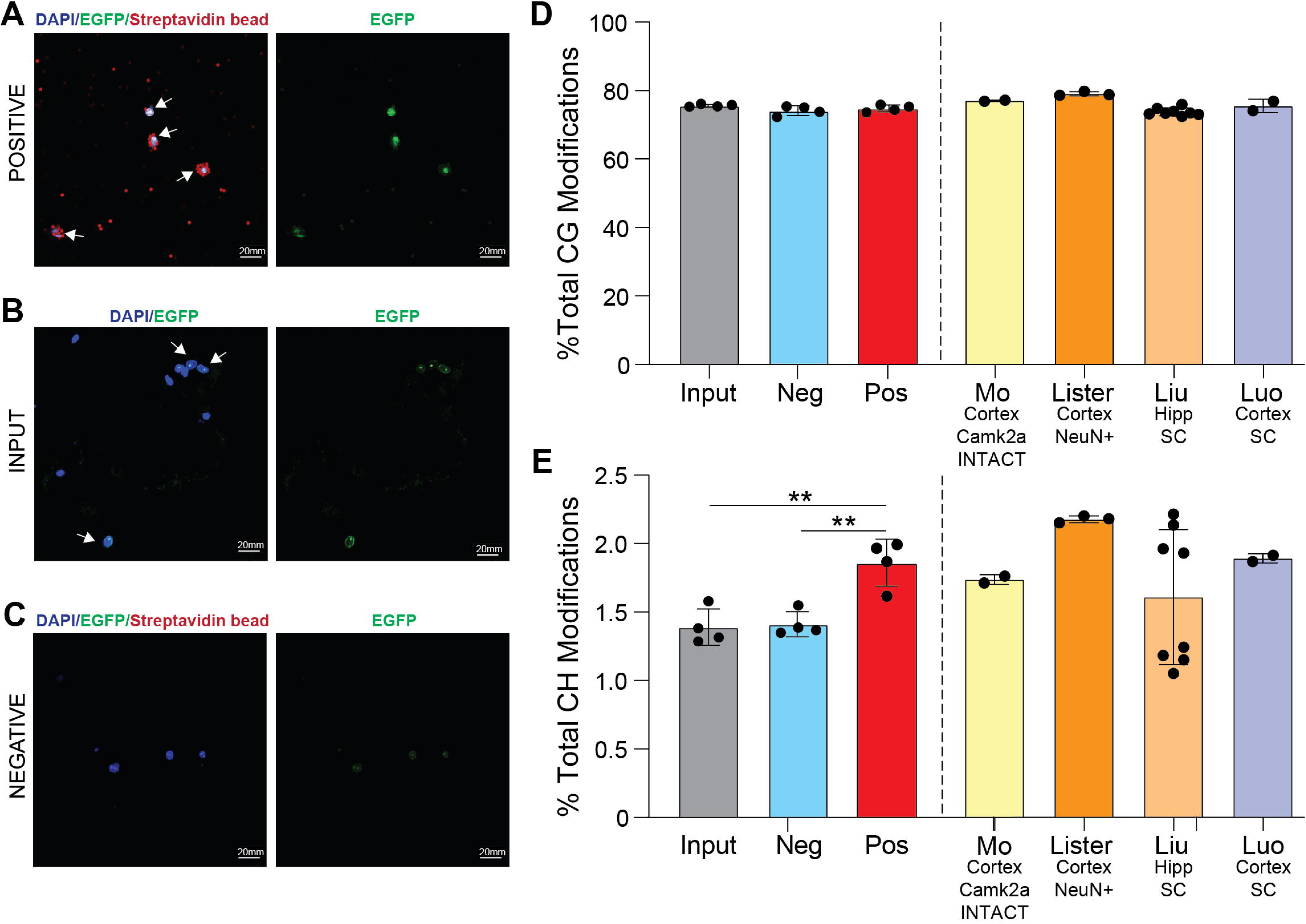
Profile of hippocampal neuronal DNA modifications by whole genome oxBS-seq. INTACT hippocampal gDNA from input, negative and positive fractions was taken for whole genome bisulfite and oxidative bisulfite sequencing. **A-D)** Total genomic levels of mCG, hmCG, mCH, and hmCH (n=4/group; one-way ANOVA with Tukey’s multiple comparisons test, *p < 0.05, **p < 0.01, ***p < 0.001). Levels of mCG were lower and hmCG and mCH were higher in the positive fraction. **E-G)** mCG, hmCG, and mCH averaged over 200 nucleotide bins from 4kb upstream, within the gene body, and 4kb downstream of neuronal marker genes in the positive fraction and input. **H-J)** Average mCG, hmCG, and mCH for positive fraction and input 4kb upstream of the TSS, within the gene body, and 4kb downstream of the TES of neuronal genes revealed lower mCG and higher hmCG and mCH in gene bodies and downstream.

The distribution of DNA modifications (mCG, hmCG, and mCH) were mapped across genic regions (Promoter, Gene Body, Downstream) of neuronal marker genes for the input and positive fraction. In the positive fraction, the intragenic/gene body and downstream regions of neuronal marker genes showed significantly lower mCG levels and significantly higher mCH and hmCG levels compared to input (**Figure 3E-J**).

### Comparison of DNA modifications across three CNS cell types

We previously validated the use of the NuTRAP construct in two mouse lines for isolation of gDNA and RNA from astrocytes and microglia [41]. To compare the DNA modification profiles between neurons, astrocytes and microglia, previously published WGoxBS sequencing data from the positive fractions of Aldh1l1-NuTRAP and Cx3cr1-NuTRAP (GSE140271) were compared to Camk2a-NuTRAP (present study, GSE228044).

The analysis revealed that microglia have a slightly but significantly lower whole genome level of total CG modifications (bisulfite conversion only, combining mCG and hmCG) than neurons and astrocytes (**Figure 4A**). When distinguishing between mCG and hmCG, neurons exhibited a shift towards higher hmCG levels and lower mCG levels compared to astrocytes and microglia (**Figure 4B**). To better understand the origin of observed DNA modification differences between cell types, we assessed the cell type-specific expression of modification regulators [DNA methyltransferases (DNMTs), Ten-eleven translocases (TETs), and thymine DNA glycosylase (TDG)]. Microglia had significantly higher DNMT (*Dnmt1*, *Dnmt3a*, and *Dnmt3b*) expression than astrocytes and microglia, aligning with microglia having the highest levels of mCG among the three cell types. Surprisingly, despite having the lowest hmCG of the three cell types assessed, microglia also express TETs (*Tet1*, *Tet2*, and *Tet3*) at a significantly higher level compared to neurons and astrocytes (**Figure 4C**). TET2 has been previously shown to regulate the microglial type I interferon-mediated inflammatory response upon LPS administration [51], pointing to a potentially dynamic role for microglial hydroxymethylation in modulating cell phenotype.

**Figure 4:**
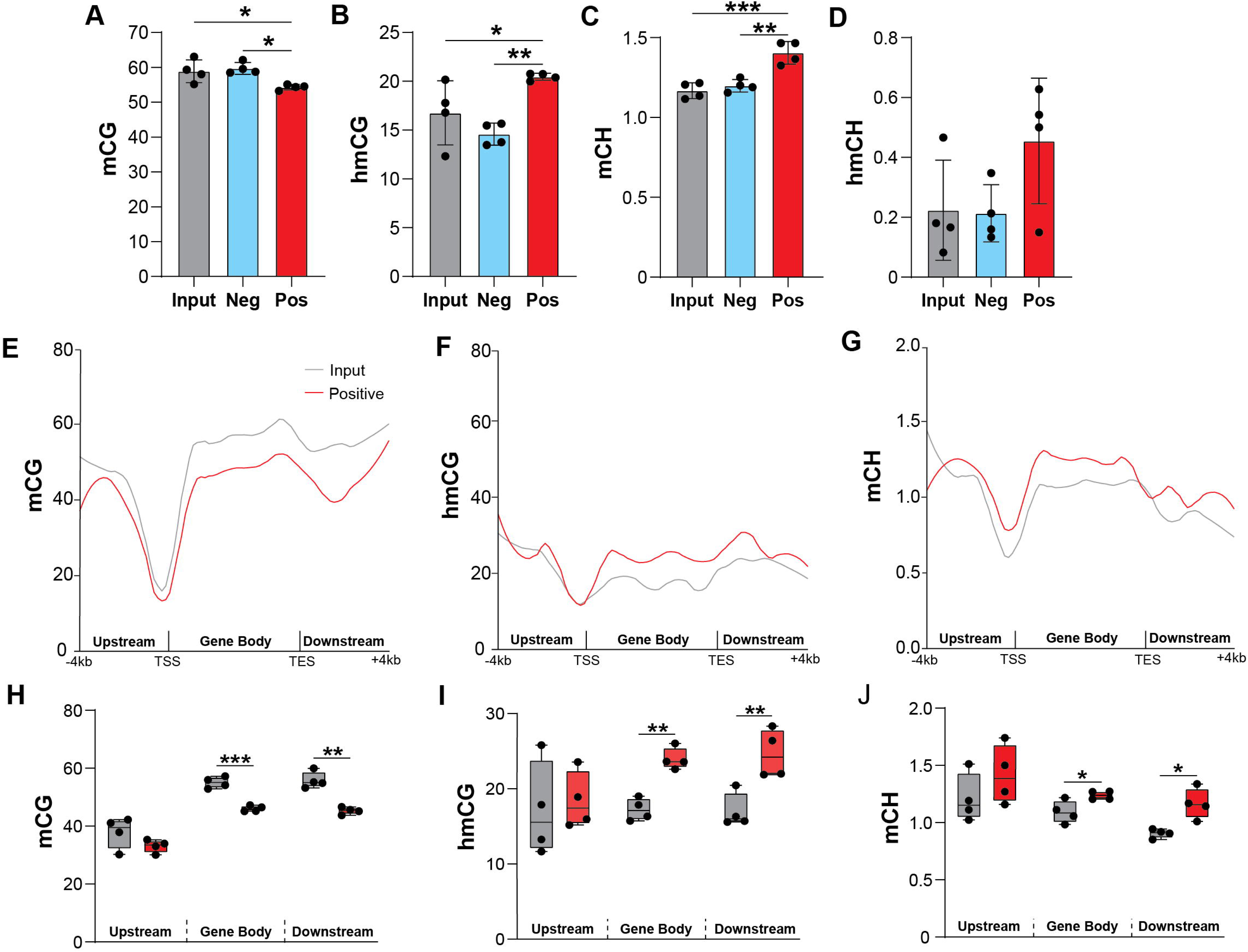
Comparison of DNA modifications across three major CNS cell types. **A)** Comparison of whole genome total CG modifications for neurons (hippocampus), astrocytes (half brain), and microglia (half brain). (n=4/group; one-way ANOVA with Tukey’s multiple comparisons test, *p < 0.05) **B)** Percentage of whole genome levels of CG, hmCG, and mCG for neurons, astrocytes, and microglia. **C)** TRAP RNA-seq expression of DNA modification regulators in neurons, astrocytes, and microglia (n=3-6/group; one-way ANOVA with Tukey’s multiple comparisons test, *p < 0.05, **p < 0.01, ***p < 0.001, ****p < 0.0001) data presented at reads per kilobase mapped. **D-F)** Whole genome, repetitive element, and non-repetitive element mCG, hmCG, and mCH levels for neurons, astrocytes, and microglia. (n=4/group; two-way ANOVA with Tukey’s multiple comparisons test, *p < 0.05, **p < 0.01, ***p < 0.001, ****p < 0.0001).

On the other hand, TDG, which mediates base-excision repair in active demethylation and single-strand break repair, was most highly expressed in neurons (**Figure 4C**). As such, the methylation and demethylation cycle may serve as a source of site-specific neuronal single-strand breaks that have been previously observed within enhancer elements [52].

Repetitive elements comprise over 50% of the genome and are thought to play an important role in neuronal differentiation and maturation [53, 54]. To determine the genomic localization of the observed cellular DNA modification differences, the levels of DNA modifications in whole genome, repeat elements, and non-repeat elements were assessed in neurons, astrocytes, and microglia. mCG levels were lower in neurons across the whole genome, repeat, and non-repeat elements compared to astrocytes and microglia (**Figure 4D**). Conversely, hmCG levels were significantly higher in neurons across repeat and non-repeat elements than astrocytes and microglia, with microglia exhibiting the lowest hmCG levels among the three cell types (**Figure 4E**). In the CH context, a similar pattern was observed across the genome and when split between repeat and non-repeat elements (**Figure 4F**).

Furthermore, when examining the split of whole genome levels into repeat and non-repeat elements, it was observed that repetitive elements had significantly higher mCG levels, while non-repetitive elements had significantly lower mCG levels compared to whole genome levels (**Figure 4D**). Conversely, there was significantly lower hmCG levels in repetitive elements compared to the whole genome levels in neurons and astrocytes, with no difference between non-repetitive elements and whole genome levels for neurons, astrocytes or microglia (**Figure 4E**).

In general, CG modification levels between cell types of repetitive and non-repetitive elements followed the pattern observed in whole genome levels. On the other hand, mCH levels were consistently the highest in neurons followed by astrocytes and then microglia, regardless of the genomic context across whole genome, non-repetitive, or repetitive elements (**Figure 4F**). Unlike CG modifications, mCH levels were observed to be nearly identical across repetitive and non-repetitive elements. These findings provide insights into the cell type-specific distribution of DNA modifications across different genomic regions, including the impact of repeat elements, and highlight the distinct epigenetic landscapes in neurons, astrocytes, and microglia.

Repetitive elements are a known source of somatic mosaicism in the brain, and their aberrant activity (particularly LINE1) is implicated in several neurological and neurodegenerative diseases [55]. We next compared the DNA modification levels between neurons, astrocytes, and microglia in specific repeat elements: long interspersed nuclear elements (LINEs), short interspersed nuclear elements (SINEs), long terminal repeats (LTRs), and simple repeats. Neurons had lower mCG levels compared to astrocytes and microglia in all analyzed repeat elements (LINEs, SINEs, LTRs, and simple repeats) (**Figure 5A**). Consistent with the whole genome levels, neurons had the highest level of hmCG and mCH within LINEs, SINEs, LTRs, and simple repeats, whereas microglia had the lowest levels of these modifications (**Figure 5B-C**). Compared to whole genome levels, mCG levels were higher within repetitive elements (LINEs, SINEs, LTRs, and simple repeats), whereas repetitive hmCG and mCH levels were lower than whole genome. The only exception to this was mCH levels within simple repeats, which were higher than whole genome levels (**Figure 5A-C**). Additionally, simple repeats use more mCH and less mCG compared to other specific repeat elements analyzed (**Figure 5A,C**). Generally, the modification patterns between specific repeat elements followed the patterns observed at the whole genome level. However, there were differences in the absolute levels of DNA modifications depending on the specific repeat element analyzed.

**Figure 5:**
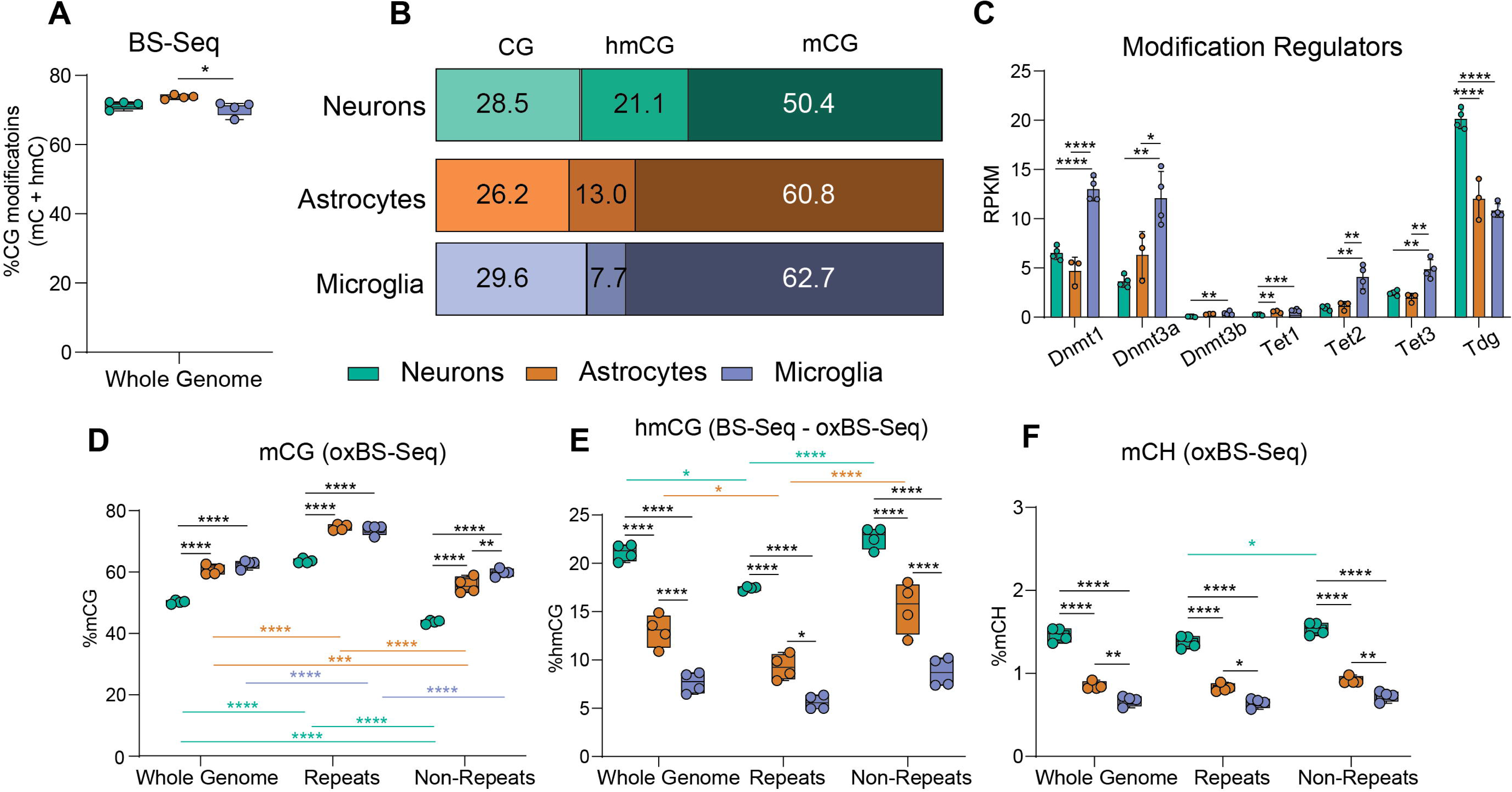
Repeat element DNA modifications in the CNS. mCG **(A)**, hmCG **(B)**, and mCH **(C)** levels of LINE, SINE, LTR, and Simple Repeat elements for neurons, astrocytes, and microglia. (n=4/group; two-way ANOVA with Tukey’s multiple comparisons test, *p < 0.05, **p < 0.01, ***p < 0.001, ****p < 0.0001).

### Differential DNA modifications are enriched at cell-type specific transcription factor motifs

To determine the genomic localization of DNA modification differences between cell types, pairwise differentially modified regions (DMRs) were identified for each DNA modification type. Differential mCG regions (DMCGRs) showed primarily comparison-specific differences, with the greatest overlap being between glia (microglia and astrocytes) and neurons (**Figure 6A,B**). DMCGRs were distributed across the genome for each comparison and ranged from -100 to 100% difference (**Figure 6C-E**).

**Figure 6:**
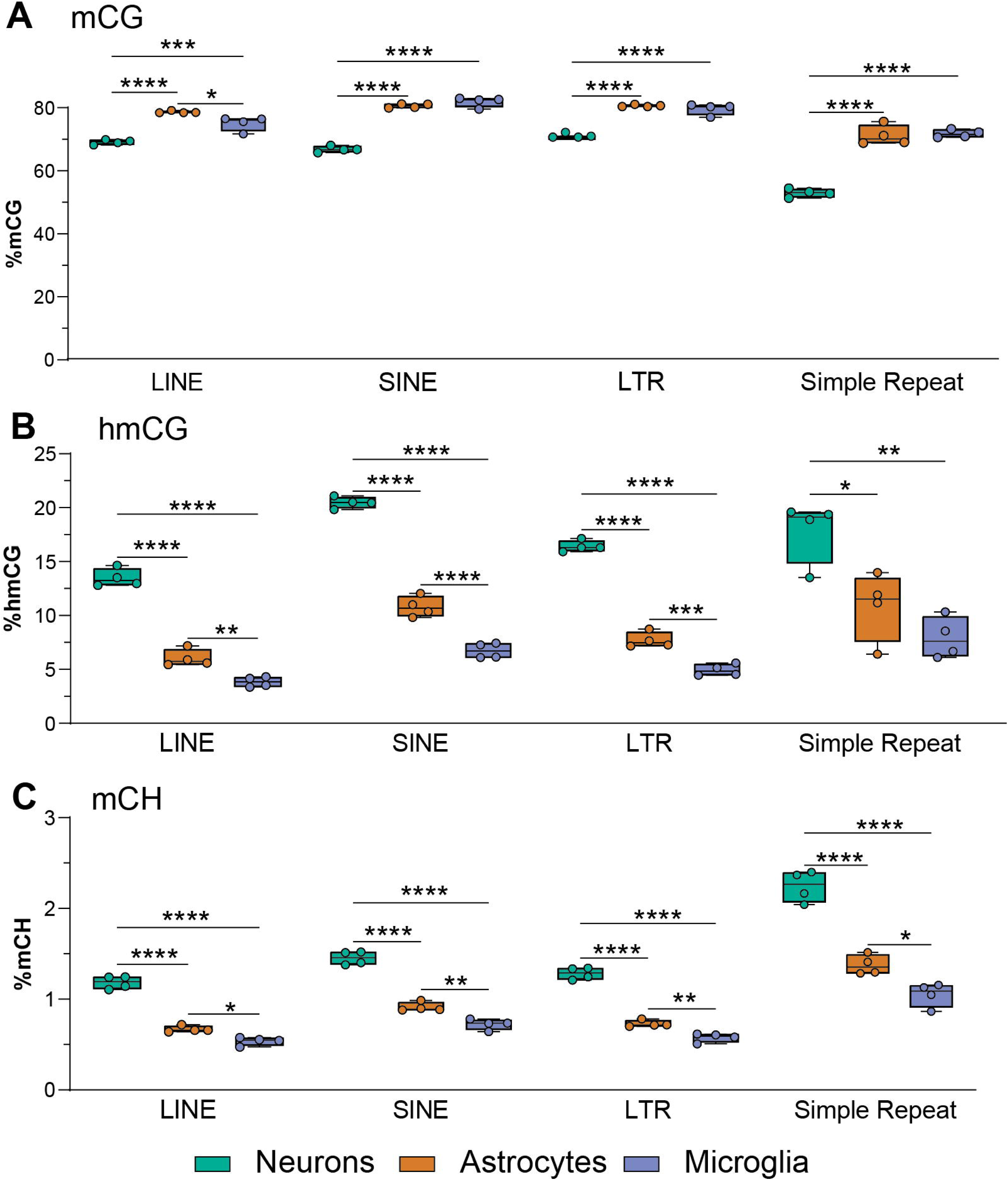
Differentially methylated CG regions. Differentially methylated CG regions (DMCGRs) were determined between cell types and overlap of hyper- **(A)** and hypo- **(B)** DMCGRs between the three comparisons. Genomic distribution and magnitude of DMCGRs for astrocytes vs neurons **(C)**, microglia vs neurons **(D)**, and astrocytes vs microglia **(E)**. Relative over- and under-representation in genic features for astrocytes vs neurons **(F)**, microglia vs neurons **(G)**, and astrocytes vs microglia **(H)**. Top enriched transcription factor binding motifs for astrocytes vs neurons **(I)**, microglia vs neurons **(J)**, and astrocytes vs microglia **(K)**.

To analyze the localization of DMCGRs within genic contexts, over- and under-representation analysis was performed (as compared to random distribution across the background). DMCGRs for all three comparisons were over-represented in gene body regions, distal promoters, and intergenic contexts, while being under-represented in proximal promoters (**Figure 6F-H**). Despite being over-represented within most cell types, astrocytic hypermethylation within distal intergenic regions compared to microglia was under-represented, demonstrating a different epigenomic patterning between individual glial cell types than between glia and neurons.

HOMER analysis was performed on DMCGRs to identify enriched motifs for each comparison (**Additional file 4**) [56], which revealed transcription factor binding motifs associated with cell type-specific functions. For instance, highly methylated astrocytic regions were enriched for HIF1b binding sites, while highly methylated neuron regions are enriched for Zic binding sites (**Figure 6I**). These transcription factors, along with their targeted genes *Npas4* and *Apoe*, have known roles in excitatory-inhibitory balance in the central nervous system [57, 58], and lipid transport in cerebellar astrocytes [59, 60], respectively. As previously mentioned, mCG hypermethylation is generally associated with transcriptional repression. Thus, hypermethylation of these essential transcription factors have downstream implications for cell type-specific functions of CNS cells.

In the comparison between microglia and neurons (**Figure 6J**), hypermethylated microglial regions were found to be enriched in Lhx2 binding sites, which inhibit *Gfap* expression and promote neurogenesis in the hippocampus [61]. Hypermethylated neuronal regions were enriched in Fli1 binding sites, which are implicated in the shift from homeostatic to ramified microglia through *Spi1* and *Runx1* [62-64]. In the comparison between astrocytic and microglial DMCGRs (**Figure 6K**), hypermethylated astrocytic regions were enriched in NF1-halfsite binding sites, which has downstream regulators such as *Sp1*, *Mef2c*, and *Sall1*, all essential modulators of homeostatic microglia [65]. Microglial hypermethylation was enriched in PU.1 binding sites, and although is most well-known for its function as a master regulator of microglia, regulates the astrocytic maturation marker *Runx2* during development as well [64]. Together, motifs in DMCGRs followed the expected inverse relationship with binding sites of known cell identity-related transcription factors.

Differential hydroxymethylated CG regions (DhMCGRs) between cell types were mainly comparison-specific, with the greatest overlap observed in hypo-hydroxymethylated regions between glial cells and neurons (**Figure 7A,B**). For all three comparisons, DhMCGRs were distributed throughout the genome and range from -100 to 100% hmCG differences (**Figure 7C-E**). Analysis of the genomic distribution of DhMCGRs revealed over-representation in genic regions and distal promoter regions, with under-representation in proximal promoter regions across all comparisons (**Figure 7F-H**). Specifically, neuronal and astrocytic hyper hydroxymethylation was under-represented in distal intergenic regions, whereas over-representation was observed in microglia (**Figure 7F-H**).

**Figure 7:**
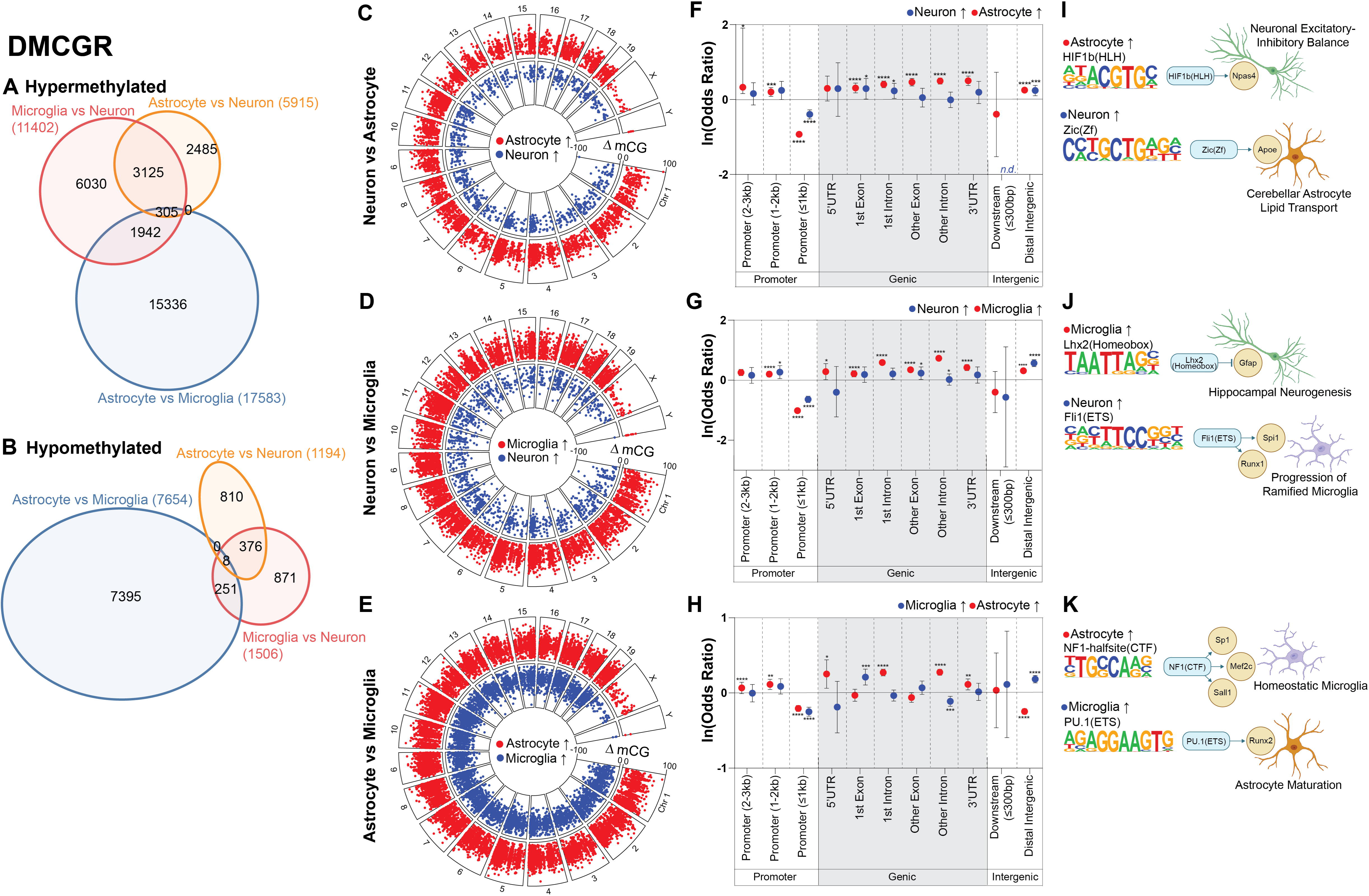
Differentially hydroxymethylated CG regions. Differentially hydroxymethylated CG regions (DhMCGRs) were determined between cell types and overlap of hyper- **(A)** and hypo- **(B)** DhMCGRs between the three comparisons. Genomic distribution and magnitude of DhMCGRs for astrocytes vs neurons **(C)**, microglia vs neurons **(D)**, and astrocytes vs microglia **(E)**. Relative over- and under-representation in genic features for astrocytes vs neurons **(F)**, microglia vs neurons **(G)**, and astrocytes vs microglia **(H)**. Top enriched transcription factor binding motifs for astrocytes vs neurons **(I)**, microglia vs neurons **(J)**, and astrocytes vs microglia **(K)**.

HOMER analysis identified enriched motifs within DhMCGRs, with corresponding cell type- specific functions (**Additional file 5**). Specifically, when comparing astrocytes and neurons (**Figure 7I**), hyper-hydroxymethylated astrocytic regions were enriched for Nr5a2 binding sites, while neuronal hyper-hydroxymethylated regions were enriched for HIF1b binding sites. Increased hydroxymethylation has been hypothesized to “functionally demethylate” specific genomic regions and positively correlate with gene expression [4]. With this in mind, hyper-hydroxymethylation of Nr5a2 (upstream of *Apoe*), in astrocytes likely induces lipid transport pathways, an essential astrocytic function [66]. Similarly, neuronal hyper-hydroxymethylation of HIF1b likely induces excitatory-inhibitory balance functions within neurons [57, 58]. Between microglia and neurons (**Figure 7J**), microglial hyper-hydroxymethylation of PU.1:IRF8 binding sites is likely involved in microglial activation programming [67]. When comparing astrocytes and microglia (**Figure 7K**), astrocytic hyper-hydroxymethylation was enriched in HIF1b binding sites, which may be needed for the central regulation of oxygen sensing, an important function of astrocytes [68]. Microglial hyper-hydroxymethylation was enriched in Six2 binding sites, which regulates an anti-inflammatory phenotype in microglia through *Gdnf* and *Il4* [69]. Taken together, the consistency of hyper-hydroxymethylated transcription factor binding motifs with specific cell type implications further bolster the hypothesis that hydroxymethylation serves to “functionally demethylate” specific regions of the genome that are needed for cellular identity.

Differentially methylated CH regions (DMCHRs) were predominantly comparison-specific, with the largest overlap observed between the microglia vs neuron and microglia vs astrocyte comparisons (**Figure 8A**). Additionally, the majority of DMCHRs were shared between glial comparisons with neurons (**Figure 8B**). DMCHRs, as with the other modification types analyzed, were distributed across the genome and also varied from -100 to 100% methylation differences, however their magnitude tended to be smaller than CG modification differences (**Figure 8C-E**).

**Figure 8:**
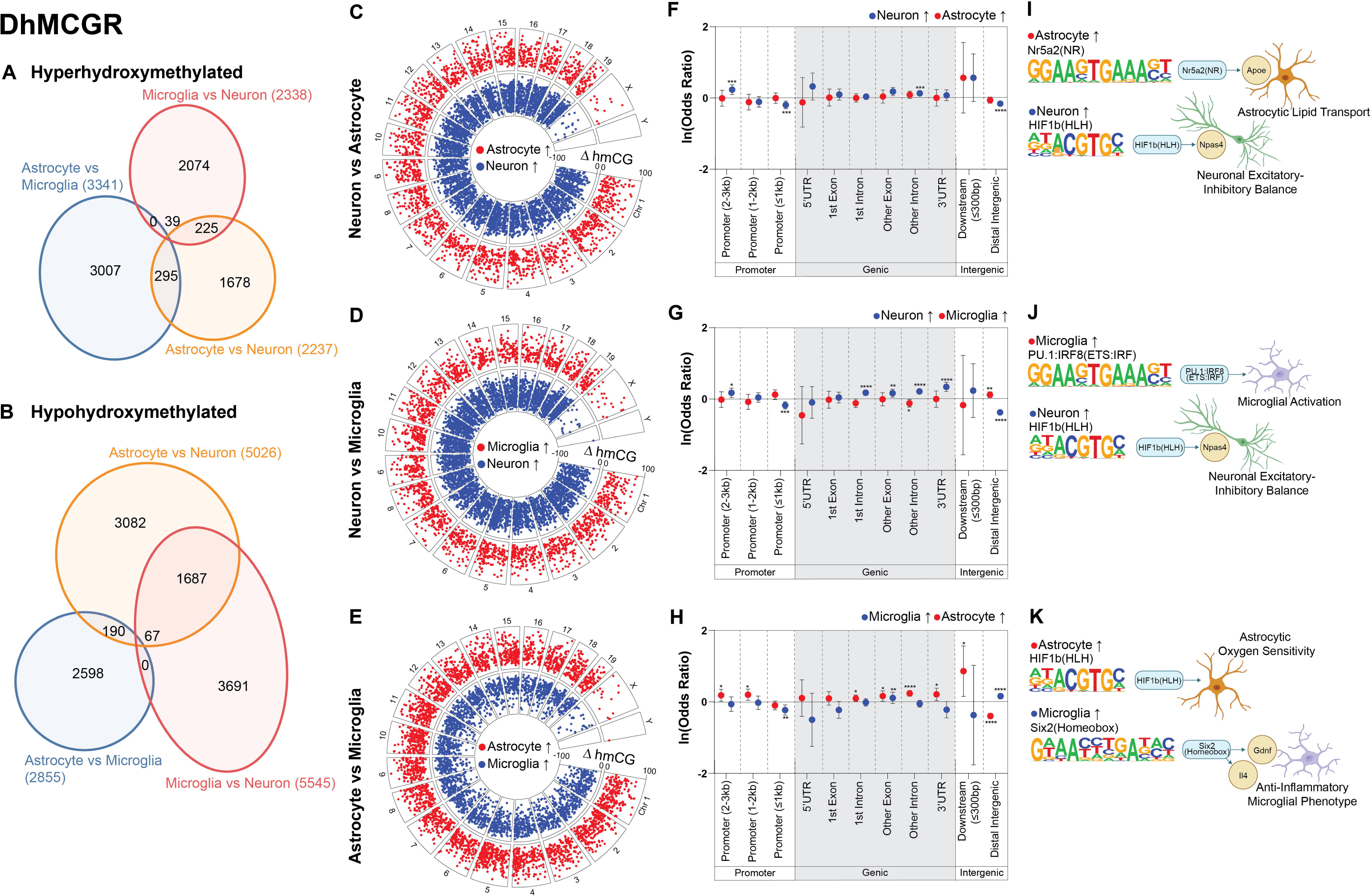
Differentially methylated CH regions. Differentially methylated CH regions (DMCHRs) were determined between cell types and overlap of hyper- **(A)** and hypo- **(B)** DMCHRs between the three comparisons. Genomic distribution and magnitude of DMCHRs for astrocytes vs neurons **(C)**, microglia vs neurons **(D)**, and astrocytes vs microglia **(E)**. Relative over- and under-representation in genic features for astrocytes vs neurons **(F)**, microglia vs neurons **(G)**, and astrocytes vs microglia **(H)**. Top enriched transcription factor binding motifs for astrocytes vs neurons **(I)**, microglia vs neurons **(J)**, and astrocytes vs microglia **(K)**.

Within all cell type comparisons, DMCHRs were over-represented in first exons and first introns, as well as distal intergenic regions (**Figure 8F-H**). Conversely, promoter regions and later gene body regions were under-represented for DMCHRs (**Figure 8F-H**). HOMER analysis identified enriched transcription factor binding sites within DMCHRs, which were associated with cell-type specific functions (**Additional file 6**). mCH has been shown to have an inverse relationship with expression [2], and this is demonstrated by the cell type-specific downstream functions of enriched binding sites in DMCHRs. In particular, between astrocytes and neurons, high astrocytic mCH regions were enriched for Klf14 binding sites, and high neuronal mCH regions were enriched in Klf10 binding sites (**Figure 8I**). Klf14 is upstream of *Pgc1a*, an important regulator of glucose metabolism in neurons [70]. Thus, hypo-CH methylation of Klf14 in neurons is consistent with its activation in this cell type and hyper-methylation in astrocytes is consistent with repression. Klf10 is upstream of *Arntl*, a regulator of astrocytic activation [71] and astrocytic hypo-CH methylation of Klf10 binding sites is consistent with utilization of this transcription factor in astrocytes.

Similarly, when comparing microglia and neurons (**Figure 8J**), high microglial mCH was enriched in Klf14 binding sites [70]. Regions of high neuronal mCH were enriched within Pdx1 binding sites, and while being most well-known for its function in the pancreas, Pdx1 has recently been recognized for its necessity in immune cells like microglia [72]. In this case, Pdx1 interacts with *Il18*, which has increased expression during microglial activation [73]. Thus, neuronal hyper-methylation of Pdx1 likely serves to repress this microglial program.

Interestingly, when comparing astrocytes and microglia, Klf14 binding sites were enriched in regions of both astrocytic and microglial hyper-CH methylation (**Figure 8K**). *H2K* is a mediator of microglial activation, which can be inhibited by Klf14 [74, 75]. Klf14 also activates *Pgc1a*, which helps to facilitate fatty acid oxidation and gluconeogenesis, both important functions of astrocytes [70]. This demonstrates differential functions for the Klf14 transcription factor in multiple CNS cell types. These findings highlight the cell type-specific functions associated with differentially methylated CH regions (DMCHRs), and give further evidence that mCH contributes to gene expression regulation not only in neurons [76] but glial cells as well.

Overall, differential DNA modifications between neurons, astrocytes, and microglia are principally located within the gene body and in distal intergenic regions, not at proximal promoters as may be expected, and have enriched transcription factor binding motifs with downstream cell type-specific functions.

### Relationship of DNA modifications to gene expression is conserved across three CNS cell types

To examine the relationship between DNA modifications and gene expression in the mouse brain at a cell type-specific level, we performed analyses of positive fraction Camk2a-NuTRAP, Aldh1l1-NuTRAP, and Cx3cr1-NuTRAP paired TRAP-RNA-seq (GSE140895, present study-GSE228043) and INTACT-WGoxBS-seq (GSE140271, present study-GSE228044) data. For each cell type, genes were separated into groups of unexpressed and expressed genes, with expressed genes being further divided into high, mid, and low expressed tertiles. Average mCG, hmCG, and mCH levels were plotted 4kb upstream, within the gene body, and 4kb downstream of those genes. As gene expression is different between cell types, the composition of the individual lists is cell type-specific (**Supplemental Figure 3; Additional file 7**). Across cell types, an inverse relationship between expression and mCG was observed at the TSS, as expected, and this relationship was maintained upstream, throughout the gene body, as well as downstream of the gene body (**Figure 9A-C**). Hydroxymethylation, while having an inverse relationship with expression at the TSS, demonstrated a positive relationship with expression within the gene body of all cell types analyzed (**Figure 9D-F**). Notably, lowly expressed neuronal genes exhibited the highest level of mCH (**Figure 9G**), which has been previously described as a mechanism for fine-tuning post-developmental gene expression through MeCP2 binding [6, 48, 77]. In contrast, mCH did not vary appreciably with gene expression in astrocytes or microglia, likely due to their low genomic mCH levels (0.5-1%) (**Figure 9H,I**).

**Figure 9:**
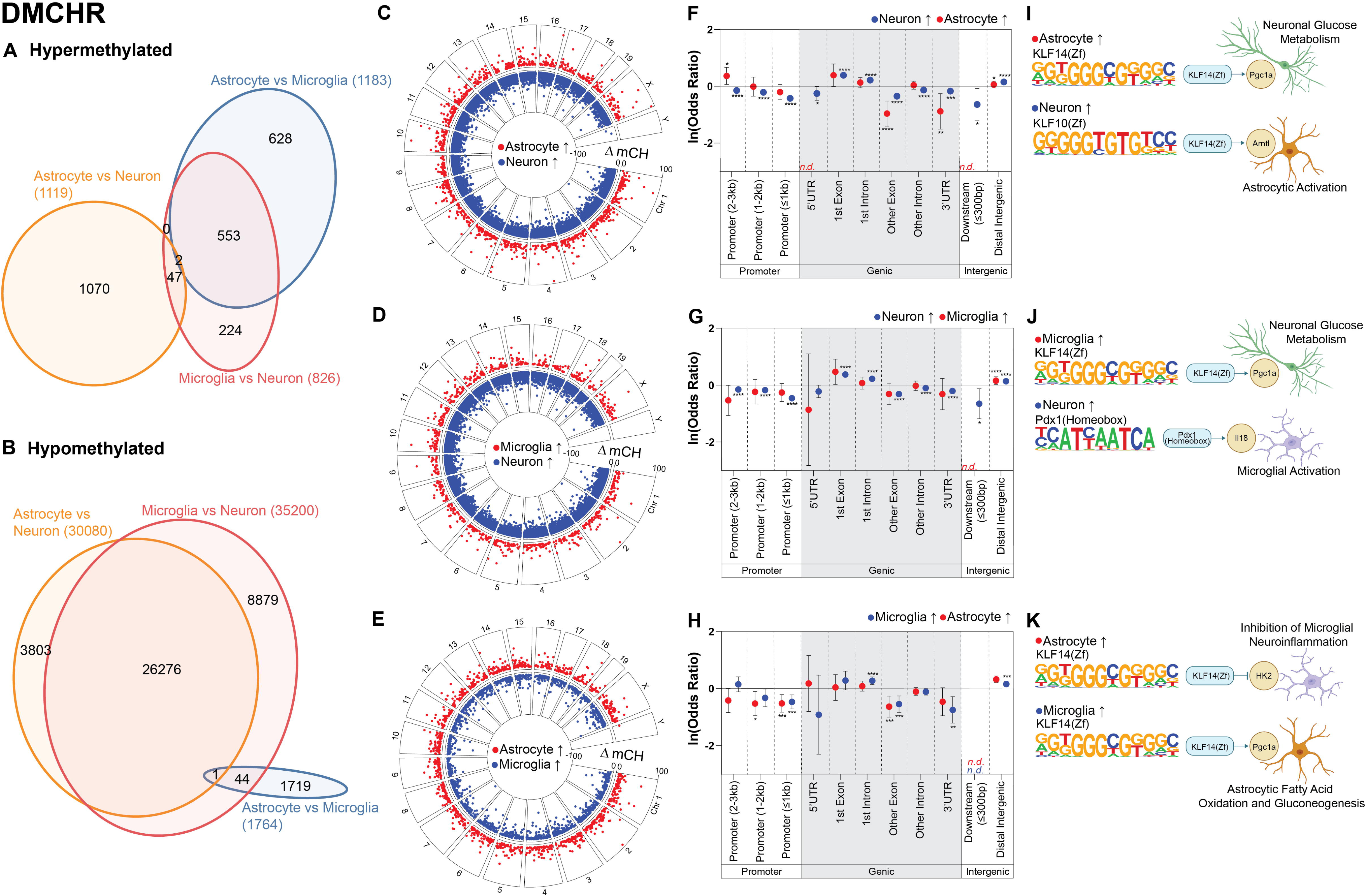
Relationship of DNA modifications to gene expression is conserved across CNS cell types. High, mid, low, and unexpressed genes were identified for each cell type from RNA-seq data. Percent mCG (**A-C**), hmCG **(D-F)** and mCH **(G-I)** averaged over 200 nucleotide bins from 4kb upstream to 4kb downstream of genes based on their expression level in each cell type.

### Integrated analysis of DNA modifications and gene expression in the CNS

To further explore the synergistic effect of different DNA modifications and cell type-specific gene expression, genome tracks of cell type-specific genes Camk2b (neuron), Atp1b2 (Acsa2; astrocyte), and Hexb (microglia) were examined. Gene expression, mCG, hmCG, and mCH for neurons, astrocytes and microglia (present study), and CpG islands (UCSC Genome Browser [78]) are displayed for each gene (raw track files can be found in **Supplemental Figure 4**). In line with previously reported functions of mCG, there was low mCG within the gene body of cell type-specific genes (**Figure 10**). There were no differences in DNA modifications between cell types at the TSS, aligning with the underrepresentation of differential modifications at proximal promoters described above. No discernable patterns were evident for hmCG or mCH (**Figure 10**), which would likely be resolved with higher sequencing depth.

**Figure 10:**
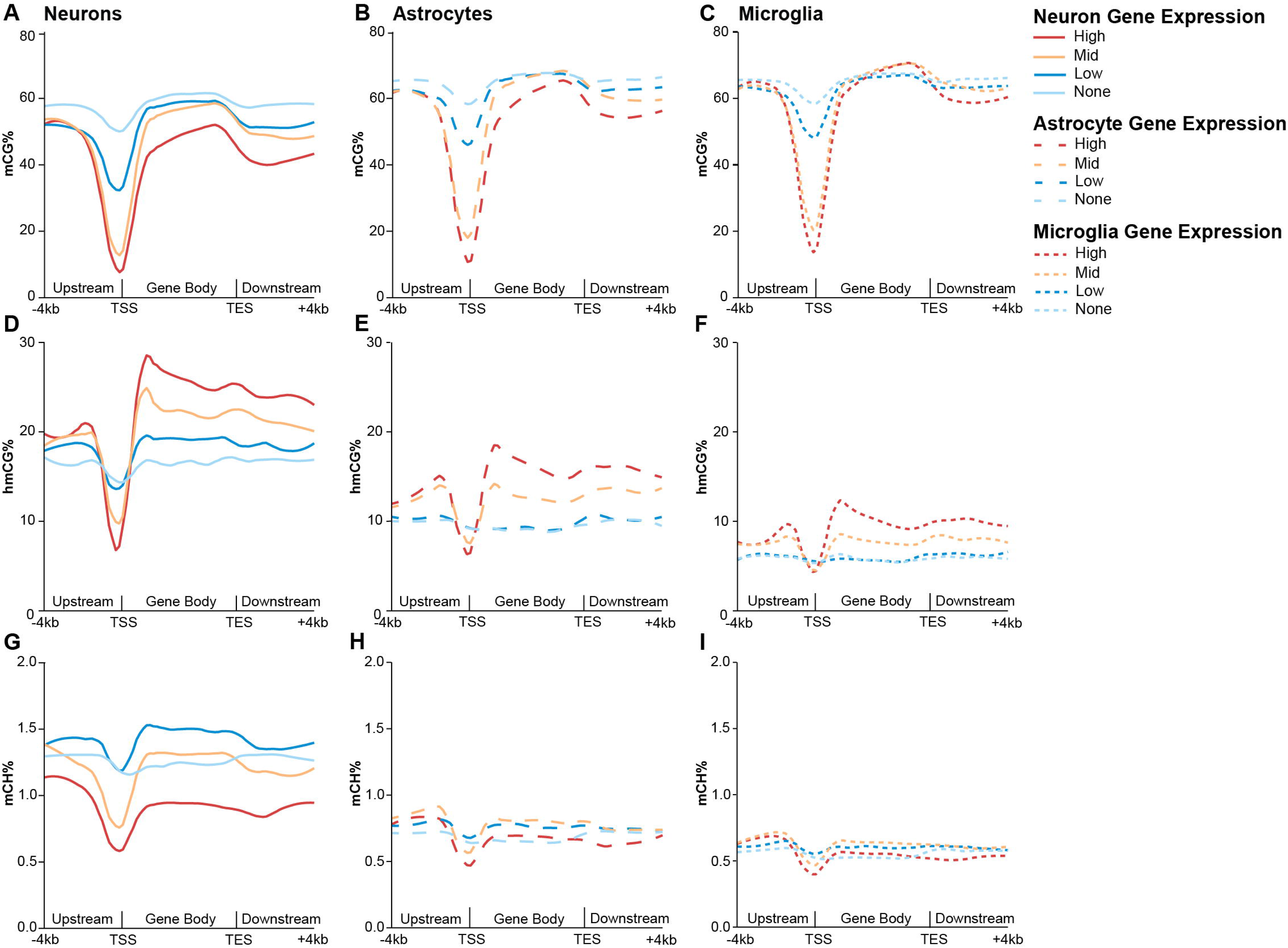
Integrated analysis of DNA modifications and gene expression in the CNS. Genome tracks for the gene body of Camk2b, Atp1b2, and Hexb with mCG, hmCG, and mCH from neurons, astrocytes and microglia compared to cell type-specific expression and CpG islands.

## Discussion

Epigenomic regulation plays a crucial role in determining cell type identity and phenotypic state [5], highlighting a need for tools to assess the neuro-epigenome at base- and cell type-specific resolution. Tissue-level analysis of the brain have provided important insights into epigenomic mechanisms regulating the genome [79-81], but the presence of mixed cell populations in whole tissue samples obscures the cell type-specificity of these mechanisms. Isolating specific cell types (i.e. via cell sorting) from the brain poses its own set of challenges, as a lack of adequate cell surface markers (particularly for neurons) and molecular changes that occur during the creation of a single cell suspension confound these types of cell type-specific analyses [82, 83]. Transgenic labeling approaches (such as RiboTag, TRAP, INTACT, and NuTRAP) provide the desired cell type specificity and avoid these activational confounds of sorting [82-84]. Adding to the growing arsenal of tools for neuro-epigenomic studies [41], here we validated a neuronal-specific Camk2a-NuTRAP model, which allows for isolation of paired DNA and RNA from hippocampal excitatory neurons for a robust cell type-specific granularity to the neuro-epigenomic regulation of gene expression.

Examining DNA modifications in specific cell types and subtypes is critical to understanding their roles in genome regulation and gene expression. However, cell type-specific DNA modification studies outside of neurons are limited, and what has been done is largely NeuN^+^ vs NeuN^-^ sorting [2, 28], which does not differentiate between glial (i.e. astrocytic, microglial, or oligodendrocytic) populations. Moreover, NeuN is a pan-neuronal marker [85], and considering that neuronal subtypes having different levels of mCG and hmCG [3], further granularity is needed here as well. This is not completely resolved even with the model presented here as Camk2a^+^ labeled neurons, while in a focused brain region (e.g., hippocampus), also represent a somewhat mixed population of pyramidal and granule neurons. However, the ability to temporally label specific neuronal populations enables more specific analysis of their epigenomic patterns. Single cell bisulfite sequencing has been conducted in the brain [49, 50], and while providing cell type specificity, is limited in its genomic coverage and does not currently have the sensitivity required to detect both mC and hmC.

The findings in this study emphasize the importance of distinguishing between methylation and hydroxymethylation when characterizing epigenome patterns and their relationship to gene expression. Bisulfite sequencing assesses does not distinguish between hmC and mC [86], and data obtained through this method are often referred to as “methylation” while actually being total modifications [87]. To circumvent these issues, antibody-based methods have been employed to assess mC and hmC, and though providing insight into overall modification levels [88-91], they do not provide the base-specificity and preferentially pull down CG dense regions of the genome making it difficult to assessment of the relationship between modifications and gene expression. Thus utilization of base-specific quantitative methods for differentiating mC versus hmC, such as oxidative bisulfite sequencing [92], enzymatic methyl-seq (EM-Seq) [93], or native reading of modifications through nanopore sequencing [94] is critical in neuroscience studies of DNA modifications.

In this study, we used validated NuTRAP models to examine DNA modifications in neurons, astrocytes and microglia. While total modification levels are similar across cell types, when split into methylation and hydroxymethylation mCG levels were highest in microglia, followed by astrocytes and then neurons. Conversely, hmCG and mCH levels were highest in neurons, followed by astrocytes and then microglia. This is in agreement with data identifying neurons as a primary source of hmCG and mCH in the brain [2-4, 95] but these findings also demonstrate the existence of hmCG and mCH in microglia and astrocytes that relates to gene expression levels. The presence of hmCH is still highly debated [2-4]. This modification is likely present in low amounts, if at all, and assessment with a direct readout of hmC such as EM-seq [96] would elucidate its presence and relationship to gene expression.

The expression of DNA modification regulators (DNMTs, TETs, TDG) did not have an obvious relationship to DNA modifications, suggesting the involvement of cell type-specific cofactors in driving these enzymes to particular genomic locations. Exploration of DNA modification regulation mechanisms in specific CNS cell typesis an important future direction for the field.

Relative modification levels across cell types in repetitive elements followed the pattern observed at the whole genome level. Repeat elements exhibited higher mCG levels and lower hmCG levels compared to non-repeat elements, indicating a strong repressive signal for repetitive elements. Transposable elements are thought to be more active during neuronal development [97-99], and the lower levels of mCG and higher levels of hmCG in mature neurons compared to astrocytes and microglia might reflect their developmental history or leave them poised for potential reactivation, although further investigation is required to elucidate this in greater detail.

The analysis of differential modification levels between cell types in this study revealed consistent trends with total modification levels, showing specific regions of hyper- or hypo-modifications for each comparison. Across comparisons, differential modifications were enriched in genic and distal intergenic regions, while being depleted within proximal promoters. This adds to a growing body of work recognizing distal regulatory regions of the genome, and not promoters, as the key regulators of cell identity [2, 3, 50, 100, 101].

Interestingly, hypomodifications were observed at proximal promoter regions, and these modifications showed an inverse correlation with expression across cell types. hmCG then demonstrated a positive association with expression in the gene body. While this relationship has been described in neurons [3, 4, 102], the relationship of hmCG to expression has not been characterized in astrocytes and microglia and provides strong evidence that hmCG is a genome regulator in glial cell types of the CNS as well as neurons. While no discernable associations between astrocytic and microglial mCH were clear, greater sequencing depth could help resolve this.

Integration of the data presented here with chromatin accessibility, histone modifications and additional genomic features by machine learning models, will contribute to a more precise understanding of the complex epigenomic regulation of gene expression that is moving beyond simplistic associations to one that is modification and context specific. While this study focused on the hippocampus, future work using the NuTRAP models and sequencing approaches that differentiate hmC from mC can be used to assess any additional CNS regions of interest. Additionally, these models and approaches can be used to examine the dynamic nature of DNA modification patterns during development, health, and neurological disorders[103, 104] in specific CNS cell types.

## Conclusions

Here, we validate a new model for studying the neuronal epigenome that circumvents the need for cell sorting, while providing greater whole genome coverage than currently available single cell techniques. While the absolute mCG, hmCG, and mCH levels across the three CNS cell types analyzed differed, the relationship of each of these modifications to gene expression is consistent across cell types. These findings demonstrate that the relationship between DNA modifications and gene expression is dependent on the genomic context and the relative modification level for that cell rather than an absolute modification level. Integration of data such as these with chromatin landscapes should reveal a more complete understanding of gene expression regulation through epigenetic mechanisms. Furthermore, gene body and intergenic region modifications, likely at enhancers, were stronger indicators of cellular identity than promoter modifications indicating that a focus on DNA modifications in proximal promoters is too simplistic.

## Methods

### Animals

All animal procedures were approved by the Institutional Care and Use Committee at the Oklahoma Medical Research Foundation (OMRF). Mice were purchased from the Jackson Laboratory (Bar Harbor, ME), bred, and housed in the animal facility at OMRF, under SPF conditions in a HEPA barrier environment. Camk2a-cre/ERT2^+/wt^ males (stock #012362) were mated with NuTRAP^flox/wt^ females (stock #029899) to generate the desired Camk2a-Cre/ERT2^+/wt^; NuTRAP^flox/wt^ (Camk2a-cre/ERT2^+^; NuTRAP^+^) progeny. DNA was extracted from ear punch samples for genotyping. Male and female mice were ∼3 months of age at the time of experiments. Euthanasia prior to tissue harvesting was carried out by cervical dislocation and decapitation. The primers used for genotyping (Integrated DNA Technologies, Coralville, IA) are included in **Additional file 1A**.

### Tamoxifen (Tam) treatment

At ∼3 months of age, mice received a daily intraperitoneal (i.p.) injection of tamoxifen (Tam) solubilized in 100% sunflower seed oil by sonication (100 mg/kg body weight, 20 mg/mL stock solution, #T5648; Millipore Sigma, St. Louis, MO) for five consecutive days. Experiments were performed 1 month after Tamoxifen administration.

### Immunohistochemistry and imaging

Brains from either Tam-induced (Tam+) or vehicle (Tam-) treated Camk2a-cre/ERT2^-^; NuTRAP^+^ or Camk2a-cre/ERT2^+^; NuTRAP^+^ mice were harvested and hemisected. Samples were fixed for 4h in 4% paraformaldehyde (PFA), embedded in 2% agarose, and vibratome-sectioned (Vibratome 3000 Sectioning System, The Vibratome Company, St. Louis, MO). Two-hundred μm-thick sagittal sections were permeated for 2h in PBS containing 3% BSA and 0.2% Triton, and processed for fluorescence immunostaining. The primary antibodies used included chicken anti-mCherry (#ab205402, 1:500, Abcam), rabbit anti-NeuN (#ab177487, 1:200, Abcam), rat anti-CD11b (#C227, 1:200, Leinco Technologies, St. Louis, MO), chicken anti-GFAP (#ab4674, 1:1000, Abcam), and hamster anti-CD31 (#2H8, 1:100, Developmental Studies Hybridoma Bank). For confocal imaging of *nuclei suspensions*, unfixed, freshly isolated nuclei were mixed with DAPI solution. Sequential imaging of nuclei was performed on a Zeiss Axiobserver Z1 Fluorescence Motorized Microscope (Carl Zeiss Microscopy, LLC, White Plains, NY) at the OMRF Imaging Core Facility. Microscope and software (Zen Black 3.1) settings were identical/similar for all samples, capture at 40X magnification. For *brain vibratome sections*, imaging was performed on an Olympus FluoView confocal laser-scanning microscope (FV1200; Olympus; Center Valley, PA) at the Dean McGee Eye Institute imaging core facility at OUHSC. Microscope and FLUOVIEW FV1000 Ver. 1.2.6.0 software (Olympus) settings were identical for samples within experiments at same magnification. The experimental format files were .oif or .oib. The final Z-stack generated was achieved at 1.22 µm step size with a total of 20 optical slices at 20X magnification (1X zoom) (Figure 1), 1.26 µm step size with a total of 22 optical slices at 20 X magnifications (1.5X zoom) (Supplemental Figure 1A,B), and 0.62 μm step size with a total of 32-50 optical slices at 40X magnification (1X zoom) (Supplemental Figure 1C-E). For all confocal images, raw files were exported as TIFF files for downstream processing and figure assembly in Adobe Photoshop V: 24.5.0 (Adobe Photoshop).

### Translating ribosome affinity purification (TRAP) and RNA extraction

The purification of cell-specific RNA from Tam-induced Camk2a-cre/ERT2^+^; NuTRAP^+^ mice (n=4) was achieved by following an established protocol [41]. One hippocampal hemisphere was minced into small pieces and homogenized in 100μL ice-cold homogenization buffer (50 mM Tris, pH 7.4; 12 mM MgCl_2_; 100 mM KCl; 1% NP-40; 1 mg/mL sodium heparin; 1 mM DTT) supplemented with 100 μg/mL cycloheximide (#C4859-1ML, Millipore Sigma), 200 units/mL RNaseOUT™ Recombinant Ribonuclease Inhibitor (#10777019; ThermoFisher), and 1X cOmplete, EDTA-free Protease Inhibitor Cocktail (#11836170001; Millipore Sigma) with a pellet pestle cordless motor (Kimble) with one 10 second pulse. 300μL ice-cold homogenization buffer was added and homogenized again with one 10 second pulse and volume brought to 1.5mL with homogenization buffer. The homogenate was transferred to a 2 mL round-bottom tube and centrifuged at 12,000 x g for 10 minutes at 4°C. After centrifugation, 100 μL of the supernatant was saved as the input. The remaining supernatant was transferred to a 2 mL round-bottom tube and incubated with 5 μg/μL of anti-GFP antibody (ab290; Abcam) at 4°C with end-over-end rotation for one hour. Dynabeads Protein G for Immunoprecipitation (#10003D; ThermoFisher) were washed three times in 1 mL ice-cold low-salt wash buffer (50 mM Tris, pH 7.5; 12 mM MgCl_2_; 100 mM KCl; 1% NP-40; 100 μg/mL cycloheximide; 1 mM DTT). After the last wash, 30μL of washed Protein-G Dynabeads were added to the homogenate/antibody mixture and incubated at 4°C with end-over-end rotation overnight. Magnetic beads were collected using a DynaMag-2 magnet and the unbound-ribosomes and associated RNA saved as the “negative” fraction (depleted). Beads were then washed three times with 1 mL of high-salt wash buffer (50 mM Tris, pH 7.5; 12 mM MgCl_2_; 300 mM KCl; 1% NP-40; 100 μg/mL cycloheximide; 2 mM DTT). Following the last wash, 350 μL of Buffer RLT (Qiagen) supplemented with 3.5 μL 2-β mercaptoethanol was added directly to the beads and incubated with mixing on a ThermoMixer (Eppendorf) for 10 minutes at room temperature. The beads were magnetically separated and the supernatant containing the target bead-bound ribosomes and associated RNA was transferred to a new tube. 350 μL of 100% ethanol was added to the tube (positive fractions: enriched in transcriptome associated to EGFP-tagged ribosomes) and then loaded onto a RNeasy MinElute column. RNA was isolated using RNeasy Mini Kit (#74104, Qiagen), according to manufacturer’s instructions. RNA was quantified with a Nanodrop 2000c spectrophotometer (ThermoFisher Scientific) and its quality assessed by HS RNA screentape with a 2200 Tapestation analyzer (Agilent Technologies).

### Quantitative PCR (qPCR)

Targeted gene expression analysis was performed with qPCR. cDNA was synthesized with the ABI High-Capacity Reverse Transcription Kit (Applied Biosystems Inc., Foster City, CA) from 25 ng of purified RNA. qPCR was performed with gene-specific primer probe fluorogenic exonuclease assays (TaqMan, Life Technologies, Waltham, MA, **Additional file 1B**) and the QuantStudio™ 12K Flex Real-Time PCR System (Applied Biosystems). Relative gene expression (RQ) was calculated with Expression Suite v 1.0.3 software using the 2^-ΔΔ^Ct analysis method with *Gapdh* as an endogenous control.

### Library construction and RNA sequencing (RNA-seq)

The NEBNext Ultra II Directional Library Prep Kit for Illumina (#NEBE7760L; New England Biolabs Inc., Ipswich, MA) was used on 25 ng of total RNA for the preparation of strand-specific sequencing libraries from input, negative, and positive fractions of each TRAP-isolated RNA sample according to manufacturer’s instructions. Briefly, polyA containing mRNA was purified using oligo-dT attached magnetic beads. mRNA was chemically fragmented and cDNA synthesized. For strand-specificity, the incorporation of dUTP instead of dTTP in the second strand cDNA synthesis does not allow amplification past this dUTP with the polymerase. Following cDNA synthesis each product underwent end repair process, the additional of a single ‘A’ base, and finally ligation of adapters. The cDNA products were further purified and enriched using PCR to make the final library for sequencing. Library sizing was performed with HS D1000 screentape (#5067-5582; Agilent Technologies) and libraries were quantified using Qubit dsDNA HS Assay Kit (ThermoFisher Scientific). The libraries for each sample were pooled at 4 nM concentration and sequenced using an Illumina NextSeq 550 (PE 75bp) at the Oklahoma Medical Research Foundation Clinical Genomics Core Facility.

### RNA-seq data analysis

Following sequencing, reads were trimmed, aligned, differential expression statistics and correlation analyses were performed in Strand NGS software package (Agilent)[105]. Reads were aligned against the Mm10 build of the mouse genome (2014.11.26). Alignment and filtering criteria included: adapter trimming, fixed 2pb trim from 5’ and 6bp from 3’ ends, a maximum number of one novel splice allowed per read, a minimum of 90% identity with the reference sequence, a maximum of 5% gap, trimming of 3’ end with Q<30. Alignment was performed directionally with Read 1 aligned in reverse and Read 2 in forward orientation. Reads were filtered based on the mapping status and only those reads that aligned normally (in the appropriate direction) were retained. Normalization was performed with the DESeq algorithm [106]. Transcripts with an average read count value >20 in at least 100% of the samples in at least one group were considered expressed at a level sufficient for quantitation and those transcripts below this level were considered not detected/not expressed and excluded, as these low levels of reads are close to background and are highly variable. For statistical analysis of differential expression, a one-way ANOVA was performed using the factor of TRAP fraction, and a Benjamini-Hochberg Multiple Testing Correction followed by Student-Newman Keuls post hoc test. For those transcripts meeting this statistical criterion, a fold change >|1.25| cutoff was used to eliminate those genes which were statistically significant but unlikely to be biologically significant and orthogonally confirmable due to their very small magnitude of change. Visualizations of hierarchical clustering and principal component analysis were performed in Strand Next Generation Analysis Software (NGS) (Version 4.0, Bangalore, India). The entirety of the sequencing data is available for download in FASTQ format from NCBI Sequence Read Archive (GSE228045). Cell type specific marker gene lists were generated from the re-analysis of lists published by McKenzie et al. [107] of immunopurified [108] and high throughput single cell data from mice [109, 110]. Published lists were filtered first by mean enrichment score of ≥3.5 and secondly to remove any genes that appeared on lists for multiple cell types. Cell population estimates within each fraction were calculated using CIBERSORTx [45], provided with raw RNA-sequencing data for each sample and cell type marker lists described above. Briefly, single-cell RNA-seq data [108] were reformatted according to the requirements of CIBERSORTx. A signature matrix was created from those data using default settings. The cellularity of each sample from TRAP RNA-seq (input, negative, and positive fractions) was imputed using the signature matrix reference and default settings. Enriched and depleted genes were imported into the Ingenuity Pathway Analysis (IPA) software (Version 01.12, Qiagen Bioinformatics) and Gene Ontology Enrichment Analysis to assess pathway/biological function enrichment, as well as identify biological processes enriched and depleted in the positive fraction compared to input.

### Isolation of nuclei tagged in specific cell types (INTACT), and gDNA and nuclear RNA extraction

Purification of viable, cell type-specific nuclei from Tam-induced Camk2a-cre/ERT2^+^; NuTRAP^+^ mouse brain tissue (n=4) was achieved by combining two previously published protocols [32, 111] with modifications as described previously [41]. One hippocampal hemisphere from the contralateral side as TRAP isolation was rinsed in ice-cold 1X PBS, minced into small pieces, and homogenized in 1 mL ice-cold nuclei EZ lysis buffer (#NUC-101, Millipore Sigma) supplemented with 1X Halt protease inhibitor cocktail (ThermoFisher Scientific) using a glass dounce tissue grinder set (#D9063; Millipore Sigma; 20 times with pestle A and 20 times with pestle B). Undissociated tissue, largely composed of blood vessels, was removed by centrifugation at 200 x g for 1.5 min at 4°C, and the supernatant containing the nuclear material filtered through a 30 μm strainer and centrifuged at 500 x g for 5 min at 4°C. The resulting nuclear pellet was resuspended in nuclei lysis EZ buffer, incubated on ice for 5 min, washed by centrifugation, and resuspended in 300 μL nuclei EZ storage buffer by gentle trituration with a micropipette. From the total resuspended pellet volume, 10% was reserved as input nuclei fraction and the rest was diluted with 1.6 mL nuclei purification buffer (NPB: 20 mM HEPES, 40 mM NaCl, 90 mM EDTA, 0.5 mM EGTA, 1X Halt protease inhibitor cocktail), and subjected to the INTACT protocol. Briefly, 30 μL of resuspended M-280 Streptavidin Dynabeads (#11205, ThermoFisher Scientific) were added into a fresh 2 mL microcentrifuge tube and washed with 1 mL of NPB using a DynaMag-2 magnet (#12321; ThermoFisher Scientific) for a total of three washes (1 min incubation/each). The washed beads were reconstituted to their initial volume (30 μL) with NPB and gently mixed with the nuclear suspension. The mixture of nuclei and magnetic beads was incubated at 4°C for 40 min under gentle rotation settings to allow the affinity binding of streptavidin beads to the cell-specific, biotinylated nuclei. After incubation, the streptavidin-bound nuclei were magnetically separated with the DynaMag-2 magnet for a period of 3 min and the unbound nuclei collected in a fresh 2 mL microcentrifuge tube, centrifuged at 4°C (1,000 x g, 3 min), resuspended in 100 μL of NPB and reserved as the negative nuclei fraction. The nuclei bound to the beads were washed in the magnet for three washes (1 min/each), resuspended in 30 μL of NPB, and reserved as the positive nuclei fraction. From each nuclear fraction [input, negative (depleted of biotinylated nuclei), and positive (enriched in biotinylated nuclei)], a 3 μL aliquot was mixed with equal volume of DAPI counterstain and used for confocal microscopy visualization and calculation of purity percentage (3-5 fields of view per sample). The AllPrep DNA/RNA Micro Kit (#80284, Qiagen, Germantown, MD) was used to extract gDNA and nuclear RNA for each sample. gDNA and nucRNA were quantified using a Nanodrop 2000c spectrophotometer (ThermoFisher Scientific) and its quality assessed by genomic DNA D1000 (#5067-5582) and High Sensitivity RNA (#5067-5579) screentapes with a 2200 Tapestation analyzer (Agilent Technologies, Santa Clara, CA).

### Library construction and oxidative bisulfite sequencing (OxBS-seq)

For each input, negative, and positive INTACT-isolated sample, 400 ng of gDNA was brought up to 50 μL volume with 1X low-EDTA TE buffer and sheared with a Covaris E220 sonicator (Covaris, Inc., Woburn, MA) to an average 200 base pare size using the following settings: intensity of 5, duty cycle of 10%, 200 cycles per burst, 2 cycles of 60 seconds, at 7°C. The size of sheared products was confirmed by capillary electrophoresis (DNA D1000 Agilent). gDNA fragments were cleaned by Agencourt bead-based purification protocol, after which gDNA was quantified (Qubit™ dsDNA ThermoFisher Scientific). Two aliquots of 200 ng gDNA fragments were prepared in a 12 μL volume to which 1 μL of spike-in control DNA (0.08 ng/μL) with known levels of specific mC, hmC, and fC at individual sites was added. End repair, ligation of methylated adaptors (#L2V11DR-BC 1-96 adaptor plate, NuGEN, Tecan Genomics, Inc., Redwood City, CA) and final repair were performed according to manufacturer’s instructions (Ovation Ultralow Methyl-Seq Library System, NuGEN). Of the two DNA aliquots per sample, one was oxidized and then bisulfite-converted and the other only bisulfite converted with the True Methyl oxBS module (NuGEN) with desulfonation and purification. qPCR was performed to determine the number of PCR cycles required for library amplification. Bisulfite and oxidative bisulfite-converted samples were both amplified for 17 cycles [95°C-2 min, N (95°C-15 s, 60°C-1 min, 72°C-30 s)]. Amplified libraries were purified with Agencourt beads and eluted in low-EDTA TE buffer. Tapestation HS D1000 was used to validate and quantify libraries. Amplified libraries were normalized to a concentration of 4 nM and pooled, denatured, and diluted to 12 pM for sequencing on the MiSeq and NovaSeq 6000 (Illumina) according to manufacturer’s guidelines with the exception of a custom sequencing primer (MetSeq Primer) that was spiked in with the Illumina Read 1 primer to a final concentration of 0.5 μM.

### OxBS-seq data analysis

Prior to alignment, paired-end reads were adaptor-trimmed and filtered using Trimmomatic 0.35. End-trimming removed leading and trailing bases with a Q-score<25, cropped 5 bases from the start of the read, dropped reads less than 30 bases long, and dropped reads with average Q-score<25. Alignment of trimmed bisulfite converted sequences was carried out using Bismark 0.16.3 with Bowtie 2 against the mouse reference genome (GRCm38/mm10). BAMs were de-duplicated with Bismark. Methylation call percentages for each CpG and non-CpG (CH) site within the genome were calculated by dividing the methylated counts over the total counts for that site in the oxidative bisulfite-converted libraries (OxBS). Genome-wide CpG and CH methylation levels were calculated separately. Hydroxymethylation levels in CpG (hmCG) and CH (hmCH) contexts were calculated by subtracting call levels from the oxidative bisulfite-converted libraries from the bisulfite-converted libraries. De-duplicated BAM files were run through methylKit in R to generate context-specific (CpG/CH) coverage text files [112]. Bisulfite conversion efficiency for C, mC, and hmC was estimated using CEGX spike-in control sequences. Untrimmed fastq files were run through CEGX QC v0.2, which output a fastqc_data.txt file containing the conversion mean for C, mC, and hmC. Analysis of methylation levels in the proximity of the promoter region was performed on a list of selected genes as follows. The R package EnrichedHeatmap was used to intersect methylation call files with genomic coordinates of gene lists [113]. Flanking regions of 4000 nucleotides were constructed upstream of the transcription start site (TSS) and downstream of the transcription end site (TES) and then split into 20 bins of 200 nucleotides each. The gene body was split into 27 equal bins, depending on the gene length. The average of each bin for all genes in the list was then plotted versus the bin number using the R package ggplot2 to give a visualization of the overall pattern of mCG, hmCG, and mCH within and around all genes contained in the gene lists [114]. Average mCG, hmCG, and mCH levels were calculated for the upstream region (-4kb to TSS), gene body (TSS to TES), and downstream region (TES to +4kb) for each gene list and biological replicate, and subjected to 2-way ANOVA statistical analysis with Sidak’s multiple comparisons correction (GSE228044). Transposable element (TE) mCG, hmCG, mCH, and hmCH was also examined. RepeatMasker BED files were obtained from the UCSC Genome Browser Table Browser (http://genome.ucsc.edu) [78]. The context-specific CpG/CH MethylKit text files were intersected with RepeatMasker BED files separated by TE type using ‘bedtools’, and percent methylation was calculated by dividing the average percent methylation at all common sites by the total number of sites. This was done for long interspersed nuclear elements (LINE), short interspersed nuclear elements (SINE), long terminal repeats (LTR), and Simple Repeats.

### DMCGR and DMCHR analysis

CpG and CH text files were read into methylKit [112] and converted into an object. The mouse genome was tiled in 1000 bp non-overlapping windows. Windows covered in all samples were retained and used for calling differentially CG/CH methylated regions with default parameters. DMRs were filtered to differences that were ≥5% between two groups with a SLIM-generated q-value less than 0.05. The methylBase and methylDiff objects were intersected to calculate percent methylation for each window passing filtering. Distribution of DMCGRs and DMCHRs within genic features and relative to the TSS was calculated using ChIPSeeker [115].

### DhMCGR analysis

To identify DhMCGRs, both BS and oxBS CpG text files were read into methylKit and converted into an object. 1000 bp non-overlapping windows covered in at least two samples were generated as above. The methylBase files generated for both BS and oxBS were read into methylKit and combined with percent methylation calculations to obtain a file containing percent methylation for each sample over windows passing filtering as described above and exported as a table. Percent methylation from BS and oxBS tiled regions was intersected using Bedtools, retaining only regions covered by both BS and oxBS. This intersected file was read back into RStudio and separated into two separate matrices containing BS and oxBS percent methylation. Hydroxymethylation over these regions was calculated by subtracting oxBS from BS for each region. DhMCGRs were filtered to differences ≥5% within each comparison (Astrocyte-Neuron, Microglia-Neuron, Astrocyte-Microglia), and assessment of the main effect of cell type was conducted using a Simple T-test (p ≤ 0.05) and manually calculated q-value ≤ 0.05 [q = min (p_i_ * N / rank_i_, q_i_ + 1)].

### DNA modification and gene expression pattern analysis

Lists of high, mid, low, and not expressed genes were generated from raw RNA-seq transcript counts of Aldh1l1-NuTRAP, Camk2a-NuTRAP, and Cx3cr1-NuTRAP positive fraction samples. Genes with zero reads for all samples were classified as not expressed, with the remaining genes being split into three equally sized lists for high, mid, and low expressed genes. The R package EnrichedHeatmap was used to intersect methylation call files with genomic coordinates of gene lists according to expression level, CGI context of the promoter, and exon. The representative plots were generated and statistical analysis performed as described for oxBS-seq analysis.

### Software usage for analysis of transcriptomic and epigenomic data

DNA modification levels across genic regions were visualized using EnrichedHeatmap in R [113]. Distribution of DMRs within genic features and relative to the TSS (Promoter (2-3kb), Promoter (1-2kb), Promoter (≥1kb), 5’UTR, 1^st^ Exon, 1^st^ Intron, Other Exon, Other Intron, 3’ UTR, Downstream (≤300bp), and Distal Intergenic) were calculated using the R package ChIPseeker [115]. Transcription factor motif analysis was performed using Homer motif analysis software (v4.10) [56], and functional interpretations were compiled using the TFLink database (https://tflink.net/) [116] and cited literature. Genomic tracks were visualized on the UCSC Genome Browser with custom tracks (http://genome.ucsc.edu) [78, 117].

## List of abbreviations

CNS: central nervous system
NuTRAP: Nuclear Tagging and Translating Ribosome Affinity Purification
Tam: tamoxifen
TRAP: Translating Ribosome Affinity Purification
INTACT: Isolation of Nuclei TAgged in specific Cell Types
WGoxBS: whole genome oxidative bisulfite sequencing
BS-seq: bisulfite sequencing
oxBS-seq: oxidative bisulfite sequencing
DNMT: DNA methyltransferase
TET: ten-eleven translocase
TDG: thymine DNA glycosylase
LINE: long interspersed nuclear element
SINE: short interspersed nuclear element
LTR: long terminal repeat
DMR: differentially modified region
DMCGR: differentially methylated CG region
DhMCGR: differentially hydroxymethylated CG region
DMCHR: differentially methylated CH region

## Declarations

### Ethical approval and consent to participate

All animal procedures were approved by the Institutional Care and Use Committee at the Oklahoma Medical Research Foundation (OMRF).

### Consent for publication

Not applicable.

### Availability of data and materials

The datasets supporting the conclusions of this article are available in the GEO repository [Camk2a-NuTRAP oxBS-seq and RNA-seq: GSE228045; Aldh1l1-NuTRAP and Cx3cr1-NuTRAP oxBS-seq and RNA-seq: GSE140271, GSE140895, GSE159106] for download in FASTQ format. Other data that support the findings of the study are available from the corresponding author (W.M.F.) upon request.

### Author’s contributions

KBT designed the work; designed and performed experiments; acquired, analyzed, and interpreted data; and wrote the manuscript. AJCE, SRO, AHM, KDP, and DRS performed experiments; acquired, analyzed, and interpreted data; and revised the manuscript. WMF designed the work; designed experiments; analyzed and interpreted data; and substantively revised the manuscript.

## Acknowledgements

The authors would like to thank the Clinical Genomics Center (OMRF), Imaging Core Facility (OMRF), Center for Biomedical Data Sciences (OMRF), Dean McGee Eye Institute (OUHSC), and Flow Cytometry and Cell Sorting Core Facility (OMRF) for assistance and instrument usage.

## Supplementary Information

**Additional file 1:** Genotyping and RT-qPCR primers. Contains the genotyping primer sequences (**A**) and RT-qPCR TaqMan gene expression assays (**B**) used.

**Additional file 2:** Cell type-specific gene lists. Contains lists of neuronal, astrocytic, microglial, oligodendrocytic, and endothelial cell-specific genes.

**Additional file 3:** Figure 2 additional information. Contains Pos vs Input fold change for each cell type specific gene (Additional file 2) from Camk2a-NuTRAP TRAP-isolated RNA-seq (**A**), CIBERSORTx results (**B**), Pos vs Input enriched genes (**C**), Pos vs Input depleted genes (**D**), GO biological processes of enriched genes (**E**), IPA functions of enriched genes (**F**), GO biological processes of depleted genes (**G**), and IPA functions of depleted genes (**H**).

**Additional file 4:** DMCGR motifs. Known HOMER motifs enriched in hyper Astrocyte vs Neuron (**A**), hypo Astrocyte vs Neuron (**B**), hyper Microglia vs Neuron (**C**), hypo Microglia vs Neuron (**D**), hyper Astrocyte vs Microglia (**E**), and hypo Astrocyte vs Microglia DMCGRs.

**Additional file 5:** DhMCGR motifs. Known HOMER motifs enriched in hyper Astrocyte vs Neuron (**A**), hypo Astrocyte vs Neuron (**B**), hyper Microglia vs Neuron (**C**), hypo Microglia vs Neuron (**D**), hyper Astrocyte vs Microglia (**E**), and hypo Astrocyte vs Microglia DhMCGRs.

**Additional file 6:** DMCHR motifs. Known HOMER motifs enriched in Astrocyte vs Neuron (A), hypo Astrocyte vs Neuron (**B**), hyper Microglia vs Neuron (**C**), hypo Microglia vs Neuron (**D**), hyper Astrocyte vs Microglia (**E**), and hypo Astrocyte vs Microglia DMCHRs.

**Additional file 7:** Genes by expression level in neurons, astrocytes, and microglia. Contains genes by expression level (high, mid, low, and non-expressed) for neurons (**A**), astrocytes (**B**), and microglia (**C**).

## Supplemental Figure Legends

**Supplemental Figure 1: Cre and Tamoxifen specificity of NuTRAP induction.** Brains were harvested from Camk2a-cre^+^; NuTRAP^+^ (Camk2a-NuTRAP) mice, treated or not with tamoxifen (Tam), for immunohistochemical analysis of NuTRAP allele recombination or for assessment of neuronal, glial, and endothelial maker expression in the context of EGFP/mCherry localization. **A-B)** Compared to counterparts from mice treated with Tam (+Tam), which exhibit robust efficiency of cre-neuronal recombination (nearly all neurons are positive for mCherry and EGFP), Camk2a-NuTRAP brains of mice not exposed to Tam (-Tam) display NuTRAP allele recombination to a subset of neurons (mCherry and EGFP expression localized to some NeuN^+^ cells). These data show a small degree of cre recombination specific to neurons independent of Tam induction (corroborating previously published observations) that is exacerbated by 5 days of systemic Tam delivery. **C)** Camk2a/NuTRAP brains show no cre recombination (EGFP or mCherry expression) in cells expressing CD11b (microglia) **D)** CD31 (endothelial), or **E)** GFAP (astrocytes). DAPI: nuclei counterstain. Scale bar: 50 μm at 20X (**A,B**), 50 μm at 40X (**C-E**).

**Supplemental Figure 2: Conversion efficiency of Camk2a-NuTRAP BS/oxBS-seq. A)** Summary of Bisulfite-sequencing (BS-Seq) and Oxidative Bisulfite-Sequencing (oxBS-Seq) techniques. Bisulfite-converted libraries are used to determine total percent modified cytosines (mC+hmC), while oxidative bisulfite-converted libraries are used to determine percent methylated cytosines (mC). hmC values are derived by subtracting oxBS from BS values on a per base basis. **B-C)** Exogenous control sequences (CEGX, Cambridge, UK) were spiked in to each sheared DNA sample (0.04% w/w) prior to oxidation and/or bisulfite conversion. Raw fastq files were read into CEGXQC v0.2 to generate summary documentation and QC reports based on the conversion efficiency of the spike-in control sequences. Conversion percentages for different cytosine modifications (C, mC, and hmC) are plotted for bisulfite-converted **(B)** and oxidative bisulfite-converted **(C)** libraries. Bisulfite-converted libraries had near complete conversion of unmodified cytosines and low over-conversion of methylated and hydroxymethylated cytosines. Oxidative bisulfite-converted libraries had high levels of conversion of unmodified and hydroxymethylated cytosines and low conversion of methylated cytosines.

**Supplemental Figure 3: Overlap of gene lists by expression level between cell types.** Venns of non-expressed **(A)**, low expressed **(B)**, mid expressed **(C)**, and high expressed **(D)** genes between neurons, astrocytes and microglia.

**Supplemental Figure 4: Raw track images from UCSC.** Raw images used to generate UCSC genome tracks from Figure 10. **A)** Camk2b **B)** Atp1b2 **C)** Hexb.

