## Supplementary figures and images for "Differential usage of DNA modifications in neurons, astrocytes, and microglia"

### Supplemental Figure 1

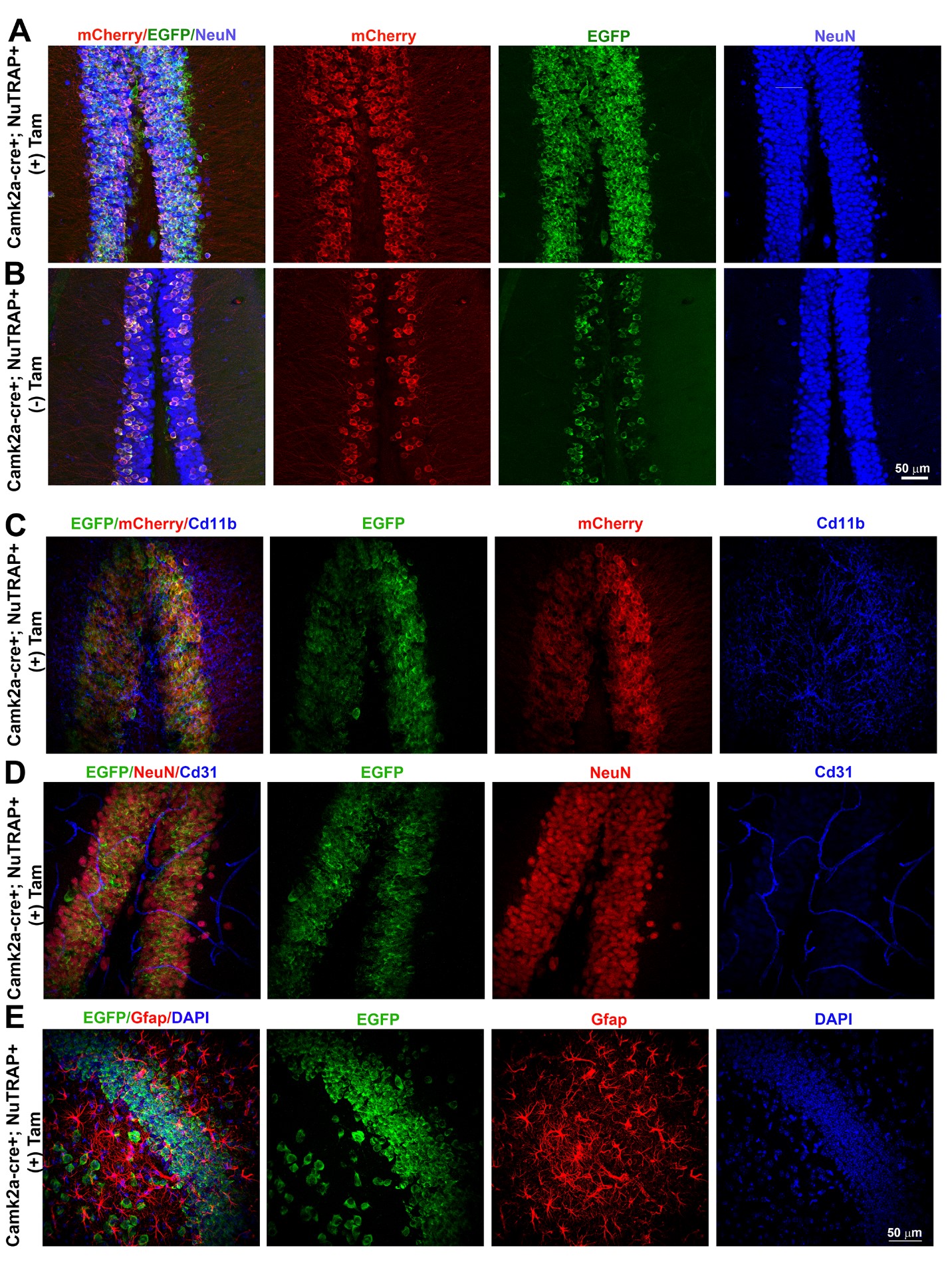

### Supplemental Figure 2

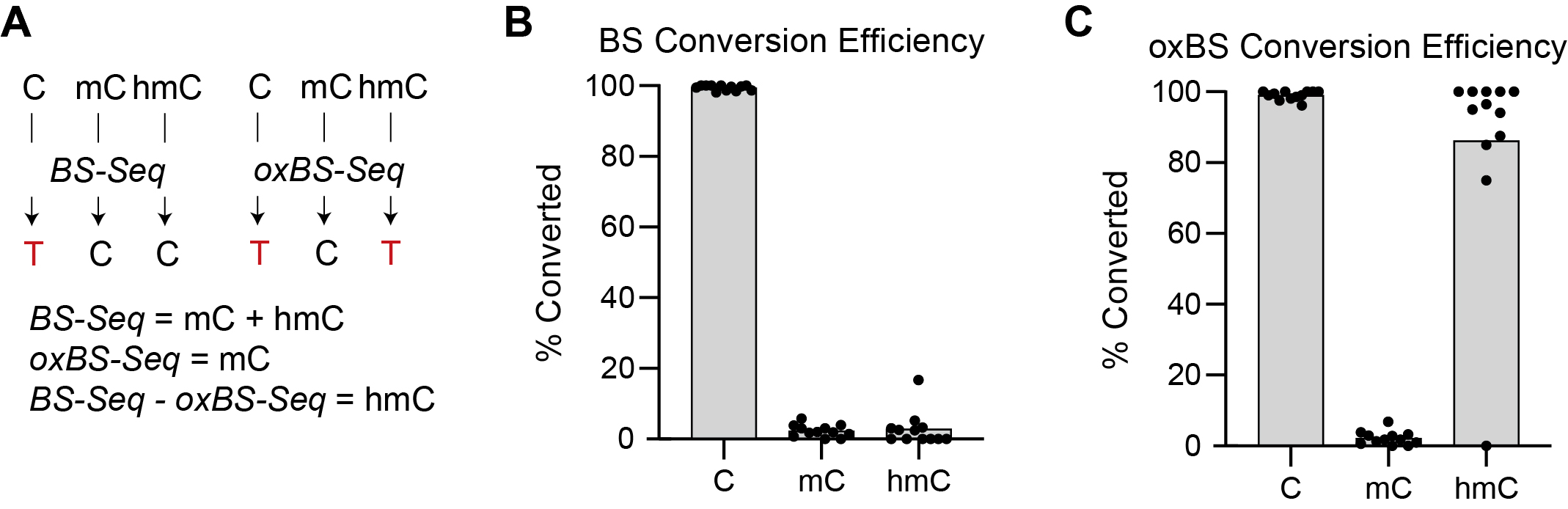

### Supplemental Figure 3

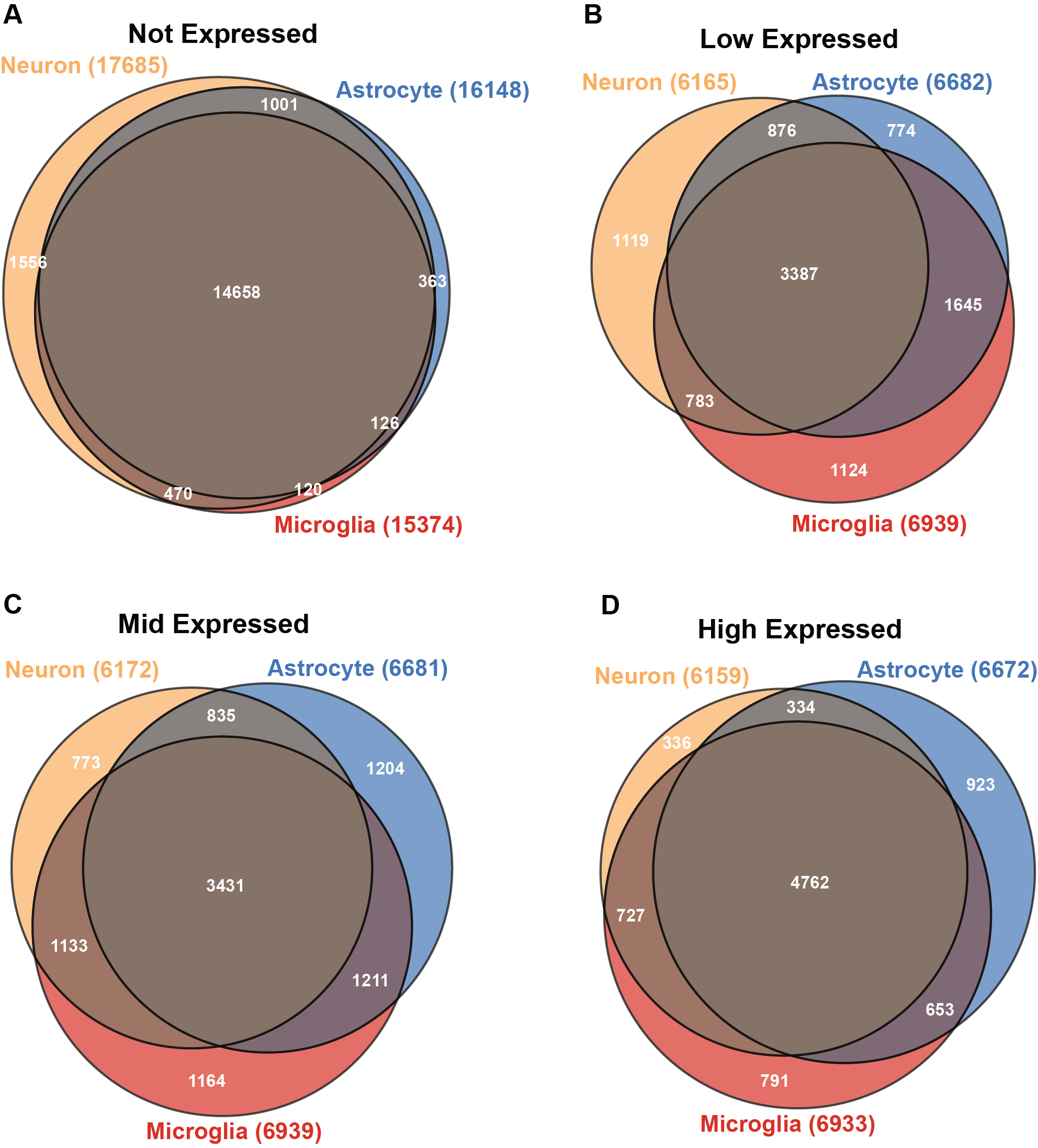

### Supplemental Figure 4

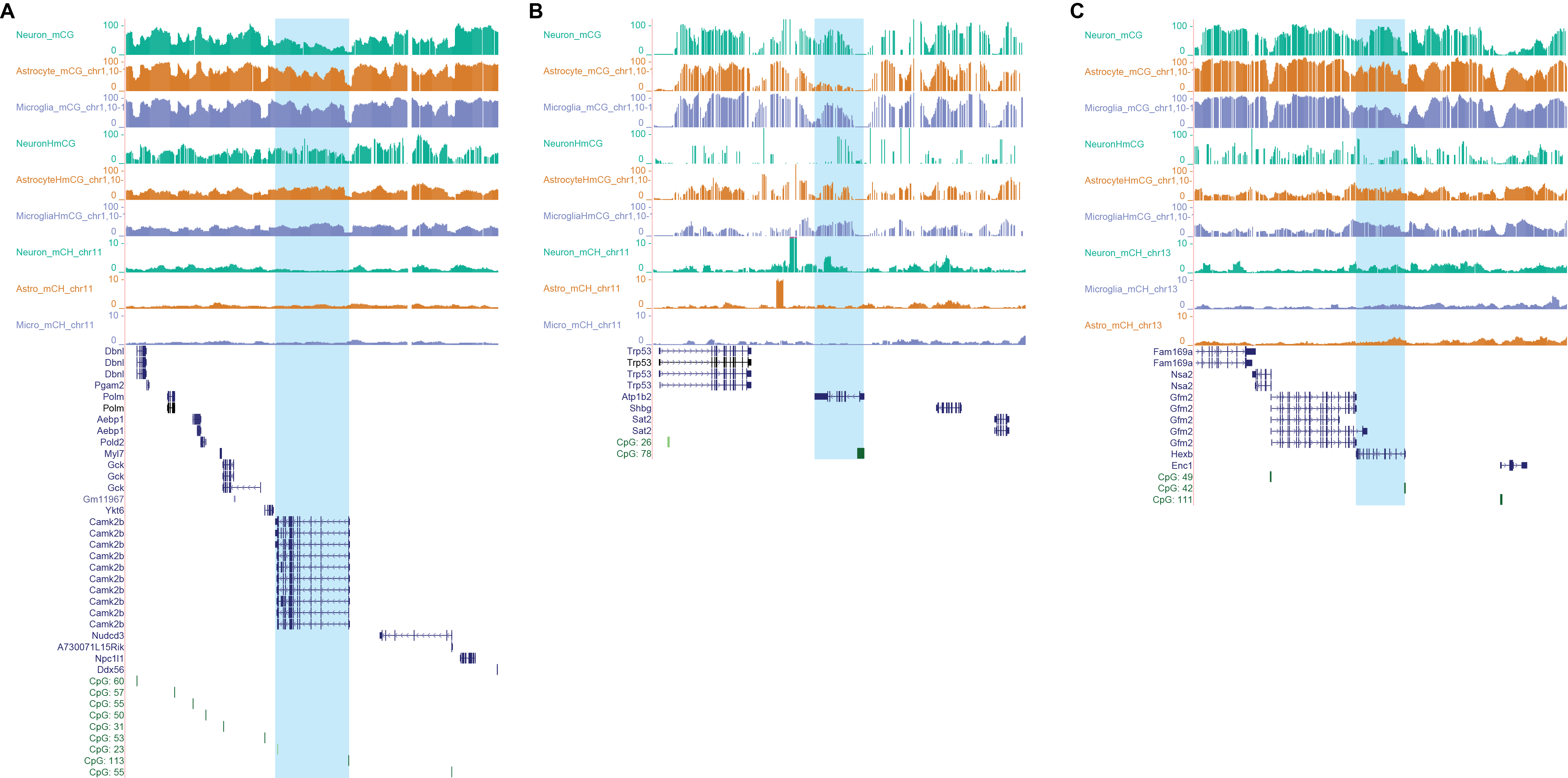
